# Transformation of Speech Sequences in Human Sensorimotor Circuits

**DOI:** 10.1101/419358

**Authors:** Kathrin Müsch, Kevin Himberger, Kean Ming Tan, Taufik A. Valiante, Christopher J. Honey

## Abstract

After we listen to a series of words, we can silently replay them in our mind. Does this mental replay involve a re-activation of our original perceptual representations? We recorded electrocorticographic (ECoG) activity across the lateral cerebral cortex as people heard and then mentally rehearsed spoken sentences. For each region, we tested whether silent rehearsal of sentences involved reactivation of sentence-specific representations established during perception or transformation to a distinct representation. In sensorimotor and premotor cortex, we observed reliable and temporally precise responses to speech; these patterns transformed to distinct sentence-specific representations during mental rehearsal. In contrast, we observed slower and less reliable responses in prefrontal and temporoparietal cortex; these higher-order representations, which were sensitive to sentence semantics, were shared across perception and rehearsal. The mental rehearsal of natural speech involves the transformation of time-resolved speech representations in sensorimotor and premotor cortex, combined with diffuse reactivation of higher-order semantic representations.

**Conflict of interest:** The authors declare no competing financial interests.

## Introduction

Immediately after hearing a series of words, we can silently replay them in our minds. What neural processes support this mental replay? Speech comprehension involves phonological, syntactic and semantic processing across widespread circuits in temporal, frontal and parietal cortex (Binder et al., 2009; Hickok and Poeppel, 2007; Pallier et al., 2011; Patterson et al., 2007; Pei et al., 2011), but experiments manipulating the load and rate of verbal rehearsal have implicated a smaller core of regions in verbal short-term memory (Fegen et al., 2015). These core areas, involved in both speech perception and production, include the posterior temporal cortex, motor and premotor areas, and the inferior frontal gyrus.

Within the regions implicated in verbal short-term memory, what kind of neural process supports the replay of recent speech? A natural hypothesis is that mental replay arises from neural replay: when we replay a series of words in our minds, the same neural populations may be activated as during the original auditory perception. This “shared representation” hypothesis is consistent with the common observation that activity patterns from perception may remain continuously active during a memory delay period or may be re-activated following periods of inactivity (Lewis-Peacock and Postle, 2008; Mongillo et al., 2008; Stokes, 2015). More generally, “reactivation” of complex sequences of perceptual input is observed during vivid imagery of those sequences (Buchsbaum et al., 2012). A shared representation for hearing and rehearsing speech would also be consistent with “mirror” models, in which the imitation of speech actions is supported by a common set of neurons across perception and production (D’Ausilio et al., 2009; Rizzolatti and Craighero, 2004).

An alternative hypothesis is that, when we silently rehearse a series of words, we employ representations that are distinct from those involved in the original auditory perception. In distinction with early observations of shared representations (Pulvermüller et al., 2006), recent intracranial and imaging studies have found that ventral sensorimotor circuits respond with distinct activity patterns during the perception and production of the same syllables (e.g., “ba”, Cheung et al., 2016; Arsenault and Buchsbaum, 2016). Moreover, widespread bilateral cortical circuits appear to “transform” between sensory and motor representations when pseudowords (e.g., “pob”) are held in mind and spoken aloud (Cogan et al., 2014). Thus, the process of mentally rehearsing an entire sentence may be supported by circuits that transform between sensory and motor representations.

We set out to determine which representations were shared and which were transformed during the perception and silent rehearsal of many seconds of natural speech. Functional magnetic resonance imaging (fMRI) studies of sentence perception and production lack the spatiotemporal resolution to map word-by-word brain dynamics at natural speech rates. Prior electrocorticography (ECoG) studies have focused on rehearsal of individual items (e.g., single syllables), lacking syntactic or semantic content and posing little demand on verbal short-term memory. Here, we used ECoG to measure time-resolved neural activity across the lateral surface of the human brain during the perception and silent rehearsal of natural spoken sentences of 5-11 words.

The processes supporting verbal short-term memory (reactivation vs. transformation) may vary according to which kind of information is being rehearsed. People are better at recalling coherent strings of words than incoherent strings of words, and this could be explained by the fact that surface features of sentences (e.g., their phonology) may be “regenerated” from abstract features (e.g., semantic and syntactics) that are most readily extracted from coherent sentences (Lombardi and Potter, 1992; Potter and Lombardi, 1990). This leads to the prediction that semantically sensitive brain regions would exhibit a “reactivation” pattern across perception and rehearsal (Bonhage et al., 2014). To test this prediction, we manipulated the internal coherence and contextual meaning of the sentences that were rehearsed.

We observed the strongest joint activation across sentence perception and silent rehearsal within the ventral sensorimotor cortex (vSMC), dorsal sensorimotor cortex (dSMC) and dorsal premotor cortex (dPMC) of the left hemisphere. Furthermore, increased activation in these areas during silent rehearsal predicted more accurate behavioral recall of the sentence content. Consistent with prior literature (e.g., Cheung et al., 2016; Glanz et al., 2018) the SMC and dPMC responded rapidly during sentence perception, encoding sub-second properties of the input. The fidelity of sensory responses in SMC and dPMC was exceeded only by the superior temporal gyrus (STG) and middle temporal gyrus (MTG). When sentences were silently rehearsed, SMC and dPMC again exhibited sentence-specific activity patterns, but the activity patterns were distinct from those observed during perception of the same sentences. Altogether, the data support a model in which “motor” circuitry (SMC and PMC) supports verbal short-term memory via a sensorimotor transformation (Cogan et al., 2014).

We also observed sentence-specific activity in anterior prefrontal cortex (aPFC) and temporoparietal cortex (TPJ). Sentence-specific activity in these areas was less temporally precise and less reliable than in sensory or motor areas. However, patterns in prefrontal areas were sensitive to the contextual meaning of the sentence being rehearsed. Moreover, the representations in these high-level areas were not transformed, but were instead shared across the perception and rehearsal of specific sentences. Activation in these higher order areas is therefore consistent with a continuous activation or reactivation process supporting verbal short-term memory.

Together, the data suggest that the sensorimotor cortex of the left hemisphere possesses both the sensory and the motor representations required to act as an audio-motor interface supporting short-term memory for natural speech sequences. “Core” speech rehearsal areas may implement a sensorimotor transformation in support of verbal short-term memory, while more distributed networks, sensitive to semantics, expressed a shared pattern of activity, bridging perception and rehearsal.

## Methods

### Participants

16 patients (11 female; 19-50 years old) were recruited from the Surgical Epilepsy Program at Toronto Western Hospital (Ontario, Canada) from the pool of all patients evaluated for neurosurgical treatment of medically refractory epilepsy. The clinical and demographic information of participants is summarized in Table S1. Prior to any experimentation, all participants provided informed consent, which was approved by the local Research Ethics Board of the University Health Network Research Ethics Board and the Internal Review Board of Johns Hopkins University. All procedures followed the Good Clinical Practice of the University Health Network, Toronto.

### Stimuli & Experimental Procedure

#### Experimental Task

Participants were asked to memorize and repeat sentences (Figure 1A-D). Each trial contained three phases: perception, silent rehearsal, and production (Figure 1A). In the perception phase, participants listened to a pair of sentences (sentence 1, S1, and sentence 2, S2). Then, in the silent rehearsal phase, participants were asked to silently rehearse S2 once verbatim in their mind, without mouthing it. Finally, in the production phase, participants were asked to vocalize the passage verbatim, at the same pace that they had heard it. Visual symbols on the screen cued participants to each phase.

**Figure 1.**
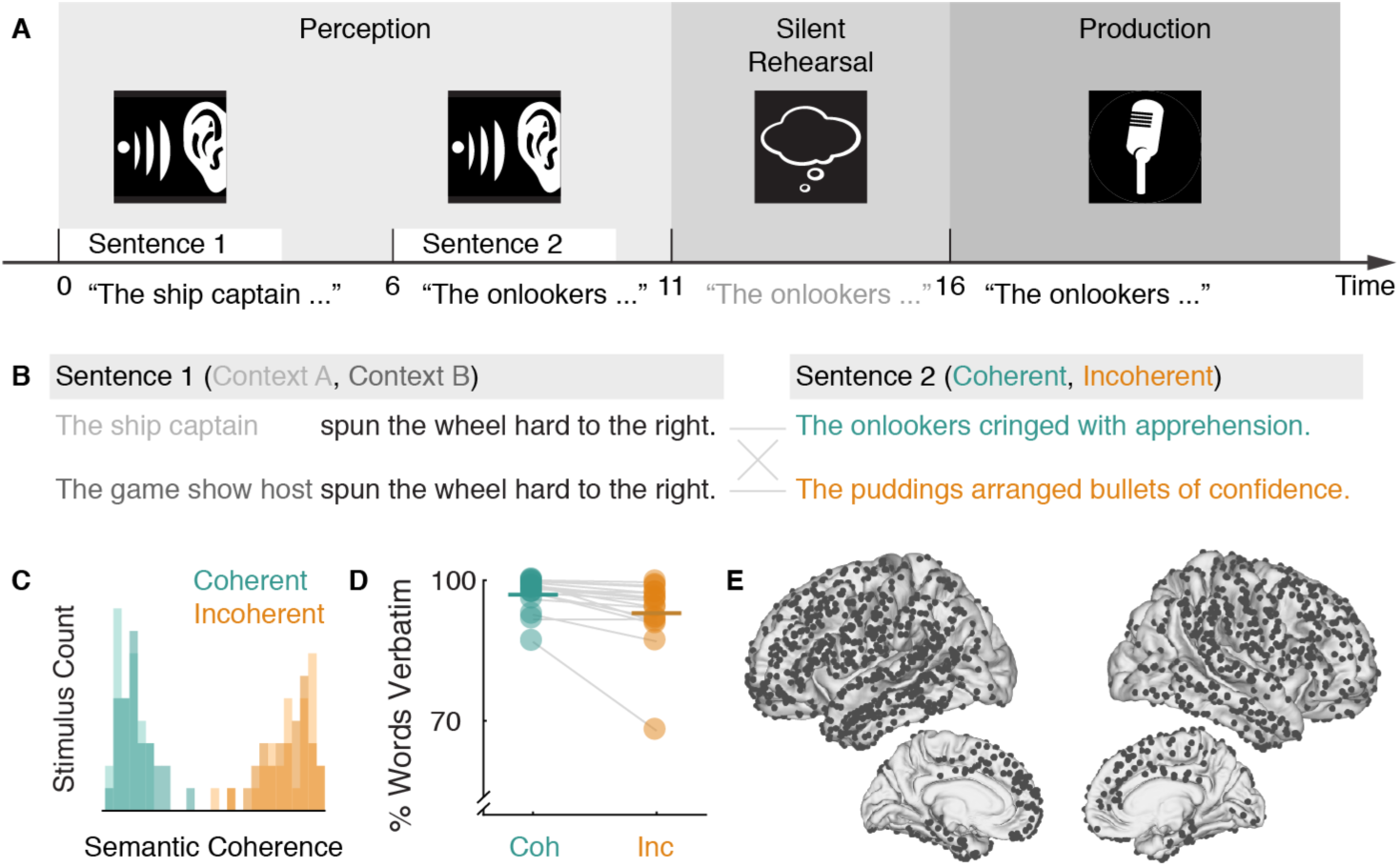
Experimental Design and Behavioral Performance. **A**, Participants first listened to two spoken sentences (sentence 1, sentence 2). They then silently rehearsed sentence 2 verbatim exactly once. Finally, they repeated sentence 2 aloud. **B**, A “stimulus group” consisted of four possible sentences: two versions of sentence 1 and two versions of sentence 2. Over the course of the entire experiment, participants would hear all four sentence 1/sentence 2 pairings. One of the sentences serving as sentence 2 was semantically coherent (teal), while the other was semantically incoherent (orange; also see Table S2). The two sentences serving as sentence 1 were both coherent but their first half differed in such a way that they would provide a very different semantic context for interpreting the coherent sentence 2. See also Table S2. **C**, Multidimensional Scaling of sentence norming data shows a clear separation of coherent and incoherent sentence combinations along the first dimension of the sentence norms. See also Table S3. **D**, Average number of words recalled verbatim for each participant when repeating coherent (Coh) and incoherent (Inc) sentences. **E**, Combined electrode placement for all 16 subjects on the lateral and medial surfaces of the Freesurfer average brain. See also Figure S1.

#### Stimulus Generation

We created 30 unique “stimulus groups” consisting of four sentences. Examples for the first sentence (S1) and second sentence (S2) are illustrated in Figure 1B, and all stimulus groups are listed in Table S2. The four sentences in a stimulus group were divided into two sentences that could serve as S1 and two sentences that could serve as S2. On a single trial, participants would hear a single pairing of S1 and S2, but across the experiment they would hear all four possible pairings. Not all participants were able to complete all 30 stimulus groups; some participants completed only 12 of the stimulus groups (set 1) while others completed an additional 18 of the groups (set 2; see Table S2).

On a given trial, the semantic context and coherence were varied, depending on which combination of S1 and S2 was presented. In half of the trials (“coherent”), S2 was semantically coherent and was a natural semantic extension of S1 (e.g., Figure 1B, top row). In the other half of trials (“incoherent”), S2 was semantically incoherent and did not have any obvious semantic relationship with S1 (Figure 1B, bottom row). For both coherent and incoherent trials, participants were asked to memorize and repeat S2 verbatim.

On coherent trials, the precise meaning of S2 depended on the contextual information presented in S1. In particular, by changing only the initial words within S1, we varied the interpretation of S2. Even the final words of S1 were shared across the two contexts. For example, the subject of the sentence in Figure 1B (“ship captain” vs. “game show host”) determines whether the “apprehension” in S2 is understood as “suspense about a prize” or “concern about danger.

For each stimulus set (set 1, set 2), we used a two-step procedure to match basic linguistic properties of the coherent and incoherent S2 sentences. First, candidate versions of semantically incoherent sentences were created using a bigram generator (http://johno.jsmf.net/knowhow/ngrams/). Second, closed-class words were replaced so that both sentences were matched for mean letter length (coherent: mean, *M* ± standard deviation, *SD* = 4.5±1.1; incoherent: *M*±*SD* 4.5±1.0; *p* = 0.795), log frequency (coherent: *M*±*SD* = 12.5±0.9; incoherent: *M*±*SD* 12.6±0.8; *p* = 0.788), number of words (coherent: *M*±*SD* = 8.1±1.5; incoherent: *M*±*SD* 8.2±1.3; *p* = 0.380) and number of syllables (coherent: *M*±*SD* = 11.0±1.3; incoherent: *M*±*SD* 11.2±1.4; *p* = 0.184). Lexical parameters were derived from the English Lexicon Project (http://elexicon.wustl.edu/default.asp). This procedure ensured that the syntax of the incoherent sentences was largely preserved.

A male speaker recorded the sentences in mono in a soundproof chamber with an MXL USB microphone (Torrance, CA). For the S1 sentences in the same stimulus group, the identical audio waveform was pasted into the shared part of each S1 sentence (e.g., “spun the wheel hard to the right” in Figure 1B). The moment when this identical acoustic waveform began in each sentence slightly differed across the two S1 sentences. Furthermore, offline processing included noise removal, cropping, volume normalization and conversion to stereo. S1 and S2 were each between 3 and 4 s in duration.

#### Norming of Sentence Pairs

An independent sample of 76 participants (40 females, 18-55+ years old) rated the 120 sentence pairs (30 stimulus groups with 4 combinations each) on eight dimensions using the online platform Amazon’s Mechanical Turk via psiTurk (https://psiturk.org/) for monetary compensation. The eight dimensions were: understandability, complexity, surprise, empathy, valence, arousal, visual vividness and auditory vividness (see Table S3 for the norming questions). Each participant rated a subset of 15 of the 120 sentence pairs on all eight dimensions (15 x 8 = 120 judgments). 2 participants were excluded because their rating data were incomplete. Following informed consent, participants were instructed to listen carefully to the audio playback of each sentence pair and make judgments using a slider on the screen. They rated each dimension on a five-point scale and were encouraged to use the full scale for their judgment. In four independent catch trials, participants had to type in the final word of the audio clip to ensure that they were paying attention to the sentence pairs. Two-factor analyses of variance were conducted on each dimension (Table S3). Coherent sentence pairs were designed to elicit a more meaningful and concrete scenario in the listeners mind. Consistent with this goal, they were easier to understand, less complex, more vivid in both visual and auditory modality, more arousing, and more empathetic than incoherent sentence pairs. Incoherent sentence pairs were more surprising than coherent sentence pairs. To visualize the similarity among the different dimensions, we applied Multidimensional Scaling on the 8-dimensional behavioral ratings of each of the 120 unique S1 and S2 combinations. The first dimension clearly separated coherent and incoherent sentence combinations (Figure 1C), confirming our manipulation of coherent and incoherent combinations.

#### Experimental Procedure

Stimuli were presented at bedside with a M-Audio M-Track Plus sound card (InMusic, Cumberland, RI) on a Dell laptop (Round Rock, TX) using the Psychophysics Toolbox 3.0.12 (http://psychtoolbox.org/) and MATLAB R2014b (Mathworks, Natick, MA) about 60 cm from their eyes. Set 1 was always presented first, while set 2 was presented if the participant was willing and able to complete the second set. All sentence pairs were repeated once, resulting in a total of 96 trials for set 1 and 144 trials for set 2. This allowed us to assess how reliable the brain responded to a given sentence. The sets were divided into eight blocks of 12 trials (set 1) or 18 trials (set 2). Each block contained all stimuli for a fixed S1 context (e.g., context 1). Half of the S2 sentences in a block were coherent and half were incoherent. The order of sentence pairs within each block was pseudorandomized such that no more than two coherent or incoherent sentences appeared in a row and that the last three stimuli were not immediately repeated in the first three trials of the next block. With two exceptions, all participants completed set 1. Participant 2 (P2) only completed one repeat for each sentence pair of set 1 (48/96). P5 completed 36/48 sentence pairs for repeat 1 and 24/48 sentence pairs for repeat 2 of set 1. P6, P7, P8, P10, P11, P13, P14, P15 and P16 completed set 2. The recordings of P14 for set 2 were discarded because of poor data quality (slow drift resulting in clipped recordings), resulting in a total of eight patients completing both sets.

To ensure that participants attended to both sentences, we presented 15 catch trials. After a completed trial the participants had to judge whether a presented picture matched with the situation described by the entire sentence pair (the combination of S1 and S2) that they had heard. Catch trials were only presented after coherent trials. On average, participants correctly responded to 80.2±3.7% (*M* ± standard error of the mean, *SEM*) of the catch trials.

### Data Acquisition & Electrode Localization

Signals were recorded from combinations of 64 contact grids, strips of 4-12 subdural platinum-iridium electrodes with 3-mm diameter and 10-mm inter-electrode spacing (PMT, Chanhassen, MN) and 4-contact Spencer depth electrodes (Ad-Tech Medical Instrument Corporation, Racine, WI). Electrode placement was solely based on clinical criteria (Figures 1E, S1). A 4-contact subgaleal strip electrode (PMT, Chanhassen, MN) electrode over the parietal midline and facing away from the brain was used for ground and reference. For data acquisition patient connectors were transferred to a separate research amplifier system (NeuroScan SynAmps2; Compumedics, Charlotte, NC). Data was recorded at 5 kHz and 0.05 Hz hardware band-pass filtering. Event markers from the stimulus presentation were sent to the SynAmps2 through a parallel port. The SynAmps2 also recorded the participant’s overt spoken response and the surrounding acoustics through a Tube MP Studio V3 (ART ProAudio, Niagara Falls, NY). Electrodes were localized with the freely available iELVIS toolbox for MATLAB (http://ielvis.pbworks.com; Groppe et al., 2017). The co-registration procedure mapped the individual structural T1 image acquired before electrode implantation on the postimplant CT image and corrected for postimplant brain shift. All electrode maps are displayed on the Freesurfer average brain.

### Data Analysis

#### Behavioral Data

Voice recordings from the task were used to manually transcribe the participant’s verbal responses exactly as uttered. For each response, we compared the transcription to the target sentence. To compute a verbatim score, we counted the words that were identical to those in the original sentence (Meltzer et al., 2016). Words did not need to be in the exactly correct order for this criterion. Words were not included if they were not part of the original sentence. Few words were recalled out of order (0.6%) or added to the sentence (2%, including metacognitive statements such as “I think, I got that wrong”). To compute an accuracy score, for each participant we computed the ratio of words recalled verbatim to the number of total words within a sentence. We computed a paired *t* test (two-sided, *α* < 0.05) to compare performance in coherent and incoherent trials.

#### Data Preprocessing

Electrodes exhibiting artifacts, epileptiform activity, excessive noise or no signal were excluded from the analysis by offline visual inspection. As the raw voltage signals of some recording sessions in a given patient exhibited different means and variance, preprocessing was done for each session separately (about 6-12 minutes of recording). Raw voltage signals were downsampled to 500 Hz. To minimize filter artifacts, all filtering was performed on continuous data. Data was high-pass filtered at 0.1 Hz (Butterworth, order 4) and line noise was removed with a bandstop filter (Butterworth) from 59.5-60.5 Hz, 119.5-120.5 Hz and 179.5-180.5. Similar to an average-rereferencing approach, the mean voltage signal across all remaining channels of a given subject was projected out from each individual channel by using linear regression. The voltage trace of each channel was z-scored, including only the trial periods for calculating the mean and standard deviation. To avoid edge artifacts, filtering was always performed on longer segments of continuous data.

Trial-level segmentation and baseline correction were performed after filtering. The filtered signal was segmented into epochs of 1 s prestimulus and 23 s poststimulus intervals, except for P1 who had a prestimulus interval of 0.5 s. We used a common trial baseline approach, in which we excluded the highest and lowest 5% of trials to minimize the effects of outlier baseline shifts: for each channel, the mean signal of the prestimulus interval was subtracted from each sample in the poststimulus interval of each trial. Bad trials were excluded from baseline correction (P6 and P10 had one bad trial each). To obtain the envelope of the acoustic data channel, we filtered the signal between 200 and 2000 Hz, applied the Hilbert transform and resampled the rectified and log-transformed signal to 500 Hz.

#### Computation of Power Time Courses

We estimated time courses of signal power using Morlet wavelets. In the frequency range 70-200 Hz, power time courses were computed separately for each frequency in steps of 5 Hz. 120 and 180 Hz were excluded as harmonics of line noise. We then took the logarithm of each power time course estimate. These estimates were then z-scored, again including only the trial periods for calculating the mean and standard deviation. Finally, we computed the mean across each of the frequency-specific z-scored time courses to yield a single “broadband” 70-200 Hz power time course.

Because of the long duration of our trials, we observed transient motor-related artifacts due to swallowing that contaminated all channels in the same frequency range as broadband power. We removed this artifact in two steps. First, we generated a temporally sparse noise regressor by identifying deviants from the median broadband power time course across all channels in a recording session. At times when the median broadband power across channels was less than three times the interquartile range from its median value, the noise regressor was set to zero; at all other times (i.e. during bursts), the noise regressor was set equal to the mean broadband power across channels. We then projected out this noise regressor from the continuous broadband power time course by taking the residual from a linear regression of the noise regressor on the broadband power signal in each channel. Before segmentation and baseline correction, broadband power time courses were smoothed with a 100 ms Hamming window.

#### Comparison of Activation in Trial vs. Baseline

Activation within trials was compared to that during baseline within each participant with a paired permutation test in sliding windows of 500 ms. The baseline window extended from −750 to −250 ms (exception: P1 had a shorter baseline period, and their window extended from - 500 to 0 ms). For each window, we computed the “activation” as the mean of the broadband power samples in that window. For each electrode, we then measured the difference between activation during the trial and the baseline period. We assessed the significance of this difference by comparing against a null distribution generated from randomly shuffling the labels (“baseline” or “trial”) 10,000 times. This procedure was computed separately for coherent and incoherent sentences. The null hypothesis of no differences between trial and baseline was rejected when the false discovery rate (FDR) across electrodes within a participant *q* < 0.01 (Benjamini and Hochberg, 1995).

#### Comparison of Correct vs. Incorrect Trials

Based on the overt memory recall in the production phase, trials were grouped into “correct” and “incorrect” bins: correct production of the sentence required an exact match to the original sentence, while all other trials were marked as incorrect. Broadband power for correct and incorrect trials were then compared with an independent-sample permutation test within each participant based on Welch’s *t*-statistic. All samples within a task phase were averaged (S1: 0-4s; gap 1: 4-6s; S2: 6-10s; gap 2: 10-11s; silent rehearsal: 11-16s; production: 16-21s). For each electrode, Welch’s *t*-statistic for the observed difference between correct and incorrect sentences was compared against a null distribution of Welch’s *t*-statistic generated from randomly shuffling the sentence labels (“correct” or “incorrect”) 10,000 times. The null hypothesis of no differences between conditions was rejected when *q* < 0.05 (FDR across electrodes within a participant, as above).

#### Comparison of Coherent vs. Incoherent Trials

Coherent and incoherent trials were compared with a paired permutation test within each participant. To rule out the possibility that activation differences could be driven by behavioral differences, i.e., incorrect recall, we only included trials on which S2 was recalled verbatim correctly or with a single small error (e.g., “a” vs. “the”). All samples within a task phase were averaged (S1: 0-4s; gap 1: 4-6s; S2: 6-10s; gap 2: 10-11s; silent rehearsal: 11-16s; production: 16-21s). For each electrode, the observed difference between coherent and incoherent sentences was compared against a null distribution generated from randomly shuffling the sentence labels (“coherent” or “incoherent”) 10,000 times. The null hypothesis of no differences between conditions was rejected when *q* < 0.05 (FDR across electrodes within a participant as above (Benjamini and Hochberg, 1995)).

#### Reliability Analysis: Correlated Component Analysis

To identify brain regions that encoded stimulus-specific information, we employed Correlated Component Analysis (Dmochowski et al., 2012). This linear decomposition technique identifies a set of weights across channels that will produce a weighted-average signal whose Pearson correlation coefficient across repeats is maximal. Intuitively, this can be thought of as assigning greater weighting to electrodes whose activity time courses are reliable across repeats of a stimulus. This technique can also be understood as similar to principal component analysis; instead of weighting electrodes such that the variance of a single repeat is maximized, this technique weights electrodes such that the correlation between two repeats is maximized, relative to the within-repeat correlation.

More formally, we define *X*_1_ and *X*_2_ as the matrices that contain the data in a given region of interest (ROI) for the first and second repeats of all stimulus groups, respectively. Each matrix has dimensions [*C* x *D*], where *C* is the number of channels in the ROI and *D* (= *T* x *S*) is the total number of data samples obtained from concatenating *T* trials with *S* samples per trial. Intuitively, the response time courses are more reliable across repeats when the diagonal values of the [*C* x *C*] covariance matrix 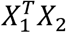 are larger. In order to find a linear combination of weights for each channel that maximizes the correlation across repeats, we solve the generalized eigenvalue problem:

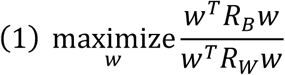

where 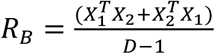 and 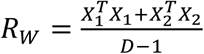 are the between-repeat covariance and the within-repeat covariance, respectively. The matrices *R*_*W*_ and *R*_*B*_ were subsequently symmetrized by taking the mean of each matrix with its transpose. To solve the generalized eigenvalue problem in a manner that is more stable, we used a shrinkage procedure to regularize the pooled within-repeat covariance matrix *R*_*W*_ (Blankertz et al., 2011). We define *R*_W_REG_=(1-*γ*)*R*_*W*_+*γλI*, where *λ* is the mean eigenvalue of *R*_*W*_, the matrix I is the identity, and γ = 0.1 is a shrinkage parameter. Thus, to estimate the weights *w* we solve the generalized eigenvalue problem with *R*_*W_REG*_ in the denominator and *R*_*B*_ in the numerator, thereby maximizing the across-repeat correlation relative to the within-repeat correlation. The [*C X 1*] generalized eigenvector with maximum generalized eigenvalue then provides the optimal weighting across the *C* channels which maximizes *ρ*.

##### Cross-Validated Correlation Across Repeats and Within Task Phases

To test reliability of responses, we employed a cross-validation procedure: weights that maximize correlation were identified in a subset of the stimulus groups, and then the reliability of the weighted signal was estimated in held-out stimulus groups. To maximize the power and stability of this procedure, we only considered data from the subjects (N=8) who completed both repeats of all 30 stimulus groups totaling 240 trials. Both correct and incorrect trials were included, because the analysis requires a pair (first and second repeat) for each trial. The electrodes from these eight subjects were pooled and grouped into nine ROIs based on the Freesurfer parcellation of the Destrieux atlas (Destrieux et al., 2010). These ROIs included STG, MTG, anterior temporal lobe (ATL), TPJ, vSMC (z<40), dSMC (z≥40), inferior frontal gyrus (IFG), aPFC (y≥23) and dPMC (y<23). Figure 2C shows the grouping of electrodes into ROIs and their average time courses for all 16 subjects. Table S4 lists the areas of the Destrieux atlas for each ROI. To be included in the pool of electrodes for this analysis, electrodes needed to show activity significantly different from baseline within at least one of the task phases (including gaps) in at least one of the conditions (coherent, incoherent).

**Figure 2.**
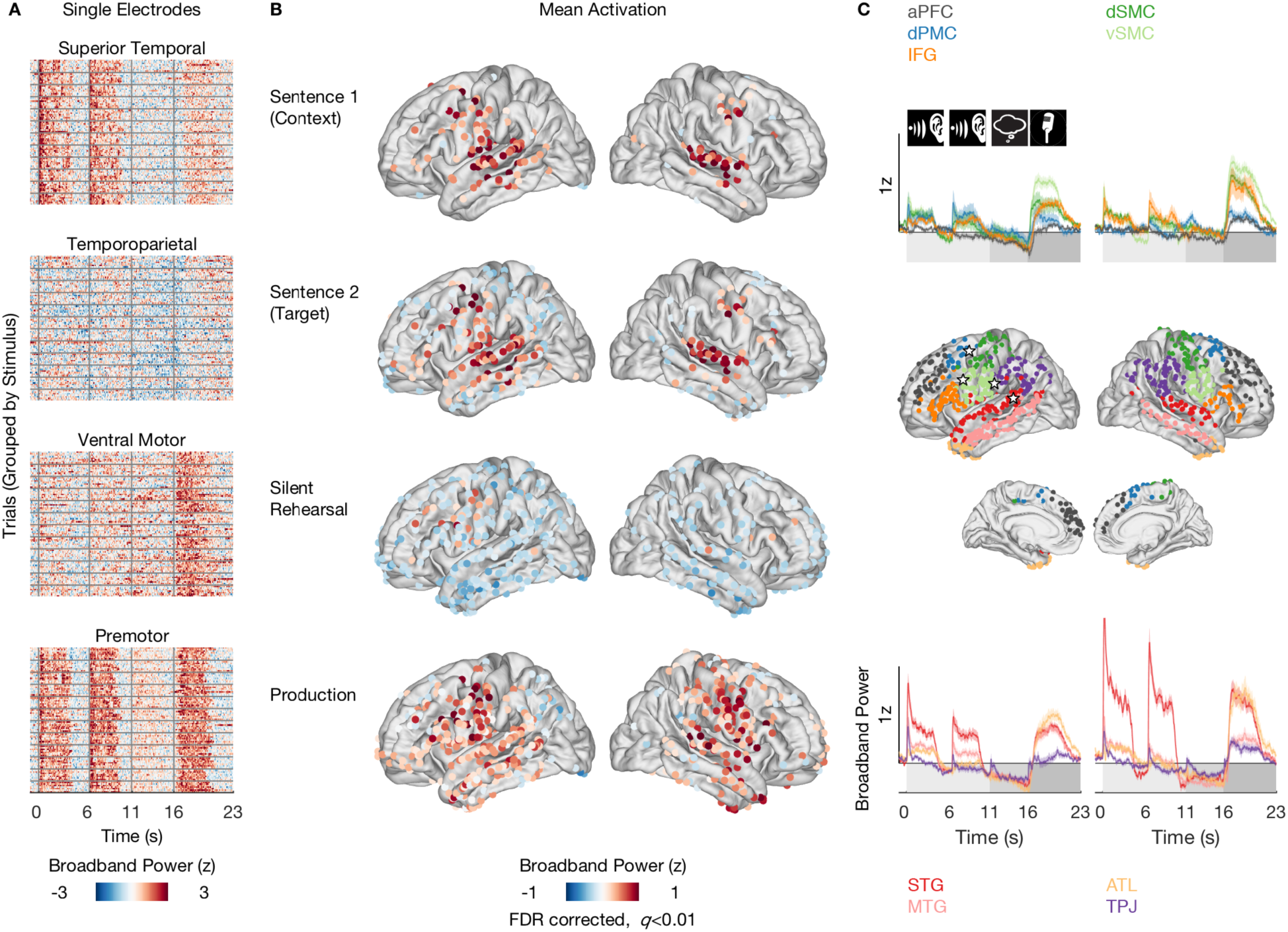
Patterns of Mean Activation During Perception, Silent Rehearsal and Production. **A**, Stacked single trial responses for example electrodes. Responses are grouped by the twelve stimulus groups shown. Broadband power changes were consistent across trials within an electrode but differed across brain regions. Electrode locations are indicated by stars (2C, middle). Onset of sentence 1, sentence 2, silent rehearsal and production are indicated by vertical gray bars. **B**, Spatial distribution of broadband power responses aggregated across participants and conditions for each task phase in the left and right hemispheres. Only electrodes activated at the level *q* < 0.01 after FDR correction are shown. See also Figure S2. **C**, Average time courses and electrode groupings within regions of interest (ROI), split by left and right hemisphere. The top panel shows the average time courses for anterior prefrontal cortex (aPFC, gray), dorsal premotor cortex (dPMC, dark blue), ventral sensorimotor cortex (vSMC, light green), dorsal sensorimotor cortex (dSMC, dark green), inferior frontal gyrus (IFG, orange). The bottom panel shows the average time courses for superior temporal gyrus (STG, red), middle temporal gyrus (MTG, pink), anterior temporal lobe (ATL, yellow) and temporoparietal junction (TPJ, purple). Onset of perception (sentence 1, sentence 2), silent rehearsal and production are indicated by icons and shading. See also Table S4.

Using a sub-sampling and cross-validation approach, weights were estimated for two thirds of the stimulus groups within a given task phase (S1, S2, silent rehearsal) and then tested on the remaining stimulus groups (Figure 5A-B). Weights were estimated separately for coherent and incoherent sentences. Thus, weight estimation sampled 40 sentences and their repeats (in-sample data: random selection of 20 stimulus groups x 2 S1 contexts x 2 repeats). Weights were then tested on the 20 held-out sentences and their repeats (out-of-sample data: the remaining 10 stimulus groups x 2 S2 contexts x 2 repeats). This procedure was repeated 1,000 times with different samples of stimulus groups to obtain a more robust estimate of repeat reliability. To avoid stimulus-specific reliability being driven by the mean response to each sentence, we normalized each trial by subtracting the mean and dividing by the standard deviation of its particular condition (sentence type, context, repeat). The normalization was performed separately for in-sample and out-of-sample data. Similar to subsampling across time, we averaged data across 200 ms bins. To feed a single time series for each channel into the Correlated Component Analysis, we concatenated all trials for a given channel.

After estimating *w* for a given ROI and task phase within in-sample data (resulting in maximum *ρ* for in-sample data), we assessed the out-of-sample reliability by measuring the correlation *ρ* in held-out data from the same task phase. In addition to measuring true repeat reliability, where each sentence is paired with its exact repetition in the held-out data, we also computed the correlation using the same weights but with trial-ordering shuffled in the matrix *X*_2_. This shuffled correlation provides a null distribution of *ρ*, against which we can compare the empirically observed, sentence-specific *ρ* values.

To assess whether the contextual manipulation (the two contexts of S1 within each stimulus group) had any effect on brain responses later in each trial, we also computed the out-of-sample correlation using the same weights, but with the responses in *X*_2_ swapped across the two versions of S1 within each stimulus group.

Overall, for each ROI we obtained 1,000 estimates of electrode weights *w* and 1,000 corresponding estimates of repeat-correlation values *ρ* for the true repeats, the null (shuffled) data, and for the repeats in which we swapped the S1 context.

##### Cross-Validated Correlation Across Repeats and Across Task Phases

In some analyses (Figures 7, 8, S4B-C), the estimated weights were also applied to held-out data from a different task phase. For those cases, we averaged all 1,000 estimates of weights *w* before we applied them to the held-out data. Since the sign of the weight is arbitrary, we first used principal component analysis to find the principal vector of weights and then multiplied all weight vectors that were opposite to that one with −1 before averaging across all weight vectors. Since the test data for this analysis is from a different task phase, it is completely separate from the data used to estimate the weights; thus, we were able to estimate the *ρ* values across task phases by subsampling from all 30 stimulus groups.

##### Statistical Assessment of Correlated Component Analysis

We assessed whether the distribution of the maximum correlation values from the 1,000 draws was stimulus-specific and significantly different from zero. To this end, we compared the true repeats of the held-out data for a given ROI against the shuffled repeats of the held-out data in a two-sided paired test for each ROI. Specifically, we computed the difference between the maximum correlation values of each fold and tested whether the 2.5^th^ and 97.5^th^ quantile contained 0. To assess an effect of the context of S1, we also compared the maximum correlation values of the true repeats with those in which the context was swapped. This test was performed one-sided because we hypothesized that the identical context elicited more reliable responses than a mismatched context. Finally, we compared coherent and incoherent sentences in a two-sample test by randomly sampling the maximum correlation values for each of the conditions 2,000 times, computing the difference, and testing whether the 2.5^th^ and 97.5^th^ quantile contained 0.

#### Cross-Temporal Analysis

To identify whether neural responses were specific to a particular moment within each stimulus, we implemented a cross-temporal analysis. This approach was inspired by cross-temporal decoding methods developed in the working memory literature (King and Dehaene, 2014). Intuitively, this analysis asks: if a particular ROI has an elevated weighted response at time *t*_1_on the first repeat, will the response also be elevated at time *t*_2_within the second repeat? If this is true, only when *t*_1_ = *t*_2_, then the neural response is specific not only to the stimulus but also specific to a particular moment within the stimulus. On the other hand, if the responses are correlated across repeats for all pairs of time-points, then the response is stimulus-specific but not timepoint-specific.

For each ROI we generated a “response time course” weighted according to the weights *w* from the Correlated Components Analysis. The response time course was generated using held-out data in each ROI both for the perception phase (listening to S2) and for the silent rehearsal phase, with the weights *w* estimated from the in-sample data for each phase. More formally, for each time point *t* within a trial and for each repeat *i* we have a matrix *Y*_*i*_(t) of the trial x channel data at that moment for that repeat. We compute a matrix of weighted time-series *Z*_*i*(t)_ = *Y*_*i*_(t)*w*, which contains the weighted neural response in that ROI for each trial. We then correlate the weighted responses of repeat 1 and repeat 2 across trials. This correlation across trials can be computed for any pair of time-points *t*_1_ and *t*_2_ leading to a time-time matrix of Pearson correlation values. If the neural response is time-point specific, then the correlations will be strong only along the diagonal of the time-time matrix, where *t*_1_ = *t*_2_. This time-time correlation procedure was repeated for the held-out data of each fold of subsampling procedure. The resulting time-time matrices were averaged across folds and sentence type (coherent, incoherent). Finally, the correlation matrices were symmetrized by taking the mean of the correlation of repeat1-to-repeat2 and the correlation of repeat2-to-repeat1.

##### De-Noising of Time-Time Correlation Maps

For visualization purposes, we performed matrix de-noising on the time-time maps using the 2-d fused lasso procedure (Tibshirani and Taylor, 2011). Briefly, this procedure estimates the original time-time maps with the constraint that many adjacent elements in the estimated time-time maps should be similar to each other. Suppose that *C* is the original time-time correlation matrix, of dimension *T* by *T*, where *T* is the number of timepoints. The 2-d fused lasso procedure solves the following optimization problem:

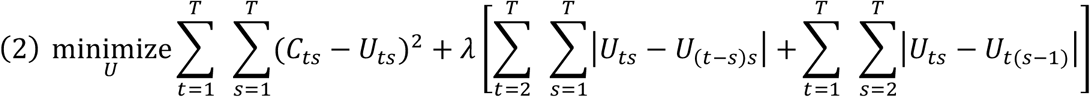

where λ is a tuning parameter that controls the number of adjacent elements that are similar to each other, and *U* is the estimated time-time map. We fit the model with several different values of λ, and the resulting matrices are plotted as heatmaps with λ = 0.03 as the best balance of smoothing and structure preservation (Figures 6, S6).

## Results

### Recall Behavior

We measured neural responses while 16 participants performed a sentence repetition task. The participants listened to a pair of sentences and were asked to mentally rehearse the second sentence and then reproduce it aloud (Figure 1A). Half of the sentences to be remembered were semantically coherent, while the other half consisted of nonsense sentences that were semantically incoherent (Figure 1B, Table S2). Participants were able to accurately reproduce both coherent and incoherent sentences in their overt recall (Figure 1D), with coherent sentences slightly better recalled than incoherent sentences (coherent: 96.9% verbatim, *SEM*: 0.9; incoherent: 93.0%, *SEM* 1.9; *t*(15) = 3.4, *p* = 0.004).

### Broadband ECoG Activity During Sentence Perception and Rehearsal

To characterize neural activity during sentence perception and rehearsal, we focused on changes in broadband power (70-200 Hz) of the local electrical field measured by subdural EcoG electrodes. Amplitude changes in this frequency band are a robust estimate of population firing (Manning et al., 2009; Miller et al., 2010; Ray and Maunsell, 2011), provide reliable responses to audiovisual stimuli across much of the lateral cortex (Honey et al., 2012) and are sensitive to speech perception and production (Cogan et al., 2014; Crone et al., 2001; Flinker et al., 2015). Aggregating data across participants resulted in dense coverage of the cortical surface in left and right hemispheres, excepting only the occipital cortex (Figure 1E).

### Patterns of Mean Activation During Perception, Silent Rehearsal and Production

Which regions of the brain were active during perception, which were active during rehearsal, and which were active during both perception and rehearsal? To quantify the overall activation patterns, we contrasted broadband power responses during each task phase against a pre-trial baseline (lateral view, Figure 2B, medial view, Figure S2A; see Methods). For S1, broadband power increased above baseline over left and right superior temporal gyrus, middle temporal gyrus, dorsal motor and premotor cortex, as well as in the left inferior frontal gyrus and anterior frontal cortex. The observation of both sensory and motor activity during passive sentence perception is consistent with prior fMRI and EcoG studies (Fedorenko et al., 2016; Honey et al., 2012; Humphries et al., 2007; Lerner et al., 2011). When participants listened to S2, which they would need to subsequently rehearse, the pattern of activation was similar, with one clear difference: a “ring” of below-baseline activity across widespread electrodes in inferior temporal, parietal and dorsal frontal regions. This ring of below-baseline activity was most pronounced in the left hemisphere, and was not observed during S1, suggesting that it was related to participants’ active attention as they prepared for the subsequent rehearsal of S2.

Silent rehearsal produced an activation pattern that was different from the perception phases: broadband activity remained above baseline only in the vicinity of the left motor and premotor cortex as well as in a very small number of lateral and inferior frontal electrodes bilaterally, and in one left posterior temporal site. Widespread electrodes across frontal, temporal and parietal cortex reduced their activity below baseline. To the extent that increases in broadband power, an estimate of population firing, index functional activation (Miller et al., 2012), these data most strongly implicate the left motor and premotor cortices in silent rehearsal.

The same patterns observed in the trial-averaged activity were present in time-resolved single-trial responses (Figure 2A). For example, a typical electrode in the superior temporal gyrus exhibited broadband power modulations that reflected auditory responses: they increased during the presentation of each sentence, showed little modulation during silent rehearsal and increased again during production, when the participant heard their own voice (Flinker et al., 2010). An example electrode in ventral motor cortex expressed slightly elevated activity during both perception and rehearsal, with the strongest modulations during production. An electrode in the premotor cortex, the area active during both perception and rehearsal, exhibited consistent single trial responses during all phases. Consistent with the observations of Cheung et al. (2016), broadband power responses in the superior temporal gyrus and premotor cortex tracked the stimulus acoustics. For example, the sample electrodes shown in Figure 2A exhibited single-trial correlations with the auditory envelope of the sentence of *r*=0.47 (superior temporal gyrus) and *r*=0.57 (premotor cortex).

To visualize the time-resolved activation patterns at a regional level, we grouped the electrodes into nine anatomically-defined ROIs (Figure 2C; see Methods). Electrodes exhibiting above or below baseline activity in at least one task phase in any stimulus condition (coherent or incoherent) were included in the regional summary. The average time courses of the ROIs reflect the overall activation pattern in Figure 2B. Although we observed above-baseline responses during sentence perception, broadband power trended towards below-baseline just until the onset of production in all ROIs (Figure 2C).

### Sensorimotor and Premotor Sites Coactive During Perception and Silent Rehearsal

Neural circuits that exhibit increased activity during both perception and silent rehearsal of a sentence are candidates for supporting the short-term memory of that sentence. Across participants, motor and premotor sites appear to be the areas most consistently involved in both perception and silent rehearsal phases of the task (Figure 2B). To spatially refine this finding, and to rule out the possibility that perceptual and rehearsal activations occurred in different subsets of participants, we measured the conjunction of sites within patients that were statistically active above or below baseline in both task phases (149 sites in total). This analysis confirmed that almost all sites active above baseline in both task phases (17 of the 20 sites) were clustered around sensorimotor and premotor cortex of the left hemisphere (Figure 3A, red electrodes). Three isolated exceptions were observed in the left posterior superior temporal cortex, the right middle superior temporal gyrus and the right inferior frontal gyrus. Sites that activated during S2 and deactivated during rehearsal (36 sites, green electrodes) clustered around superior and middle temporal cortex. Sites that were deactivated in both task phases (92 sites, blue electrodes) were mostly distributed around temporal and motor cortex (see deactivated sites during S2 in Figure 2B). Finally, there was one site in left frontal cortex that was activated during rehearsal but deactivated during perception (gray electrode). The overall results of this analysis are (i) joint perception and rehearsal activation in sensorimotor and premotor areas (see also, Glanz et al., 2018; Towle et al., 2008), and (ii) widespread suppression of rehearsal activity in a ring of surrounding temporal and parietal sites.

**Figure 3.**
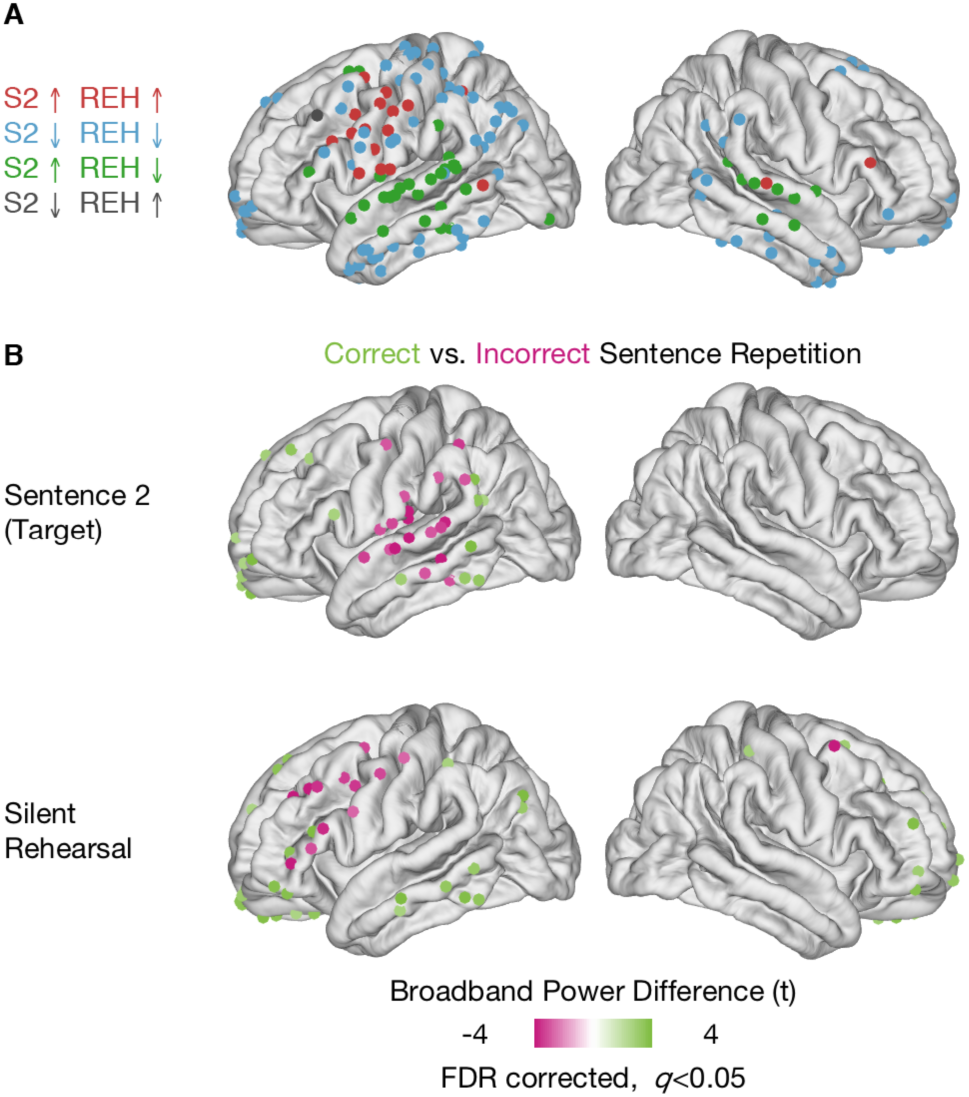
Coactivation and Memory-Dependent Behavior in Sensorimotor and Premotor Sites. **A**, Motor and premotor sites coactive during perception and silent rehearsal. Electrodes are color coded based on their combined level of activation/deactivation in the perception phase (sentence 2, S2) and silent rehearsal phase (REH). Sites that are activated above baseline in both task phases (red) cluster around sensorimotor and premotor cortex. **B**, Spatial distribution of broadband power differences for correct (light green) and incorrect sentences (pink) during sentence 2 (top) and silent rehearsal (bottom). Trials were grouped based on the overt sentence repetition behavior in the production phase. Only electrodes activated at the level *q* < 0.05 after FDR correction are shown.

### Motor and Premotor Activation Levels Predict Behavioral Accuracy

If a region encodes and maintains sentence-related information, its activity levels should be correlated with the accuracy of subsequent memory-dependent behavior. We therefore assessed the accuracy of memory-dependent behavior using the overt speech that participants produced in the production phase at the end of every trial. Trials were divided into “correct” and “incorrect” bins: correct production of the sentence required an exact match to the original sentence, while all other trials were marked as incorrect. For each electrode, we computed the difference in activation associated with correct and incorrect trials (Figure 3B).

Greater activation in the superior temporal cortex during the encoding phase (perception of S2) was predictive of less successful sentence memory (Figure 3B). The same effect was also observed in a small number of parietal and motor sites. This pattern is consistent with the role of all these areas in the initial encoding of auditory and linguistic information. To identify the regions supporting working memory, we examined activity during silent rehearsal that was predictive of accurate sentence memory (Figure 3C). When motor and premotor regions exhibited increased activation, this predicted less accurate later sentence production.

Because the behavioral associations identified in Figure 3B may be mediated by stimulus properties (e.g., sentences that are easier to remember because of their content may produce less activations) they should be interpreted with caution. Nonetheless, the results indicate that motor and premotor sites are important sites for memory-dependent behavior.

### Coherent Sentences Elicit Greater Broadband Power in Semantic Network During Perception and Rehearsal

To probe semantic selectivity, we assessed activation differences during perception and silent rehearsal when S2 was semantically coherent or incoherent (lateral view, Figure 4, medial view Figure 4). No electrodes exhibited differences between coherent and incoherent trials during S1 or gap 1. This was expected, since the nature of the trial (coherent S2, incoherent S2) is not known to the participant before the onset of S2.

**Figure 4.**
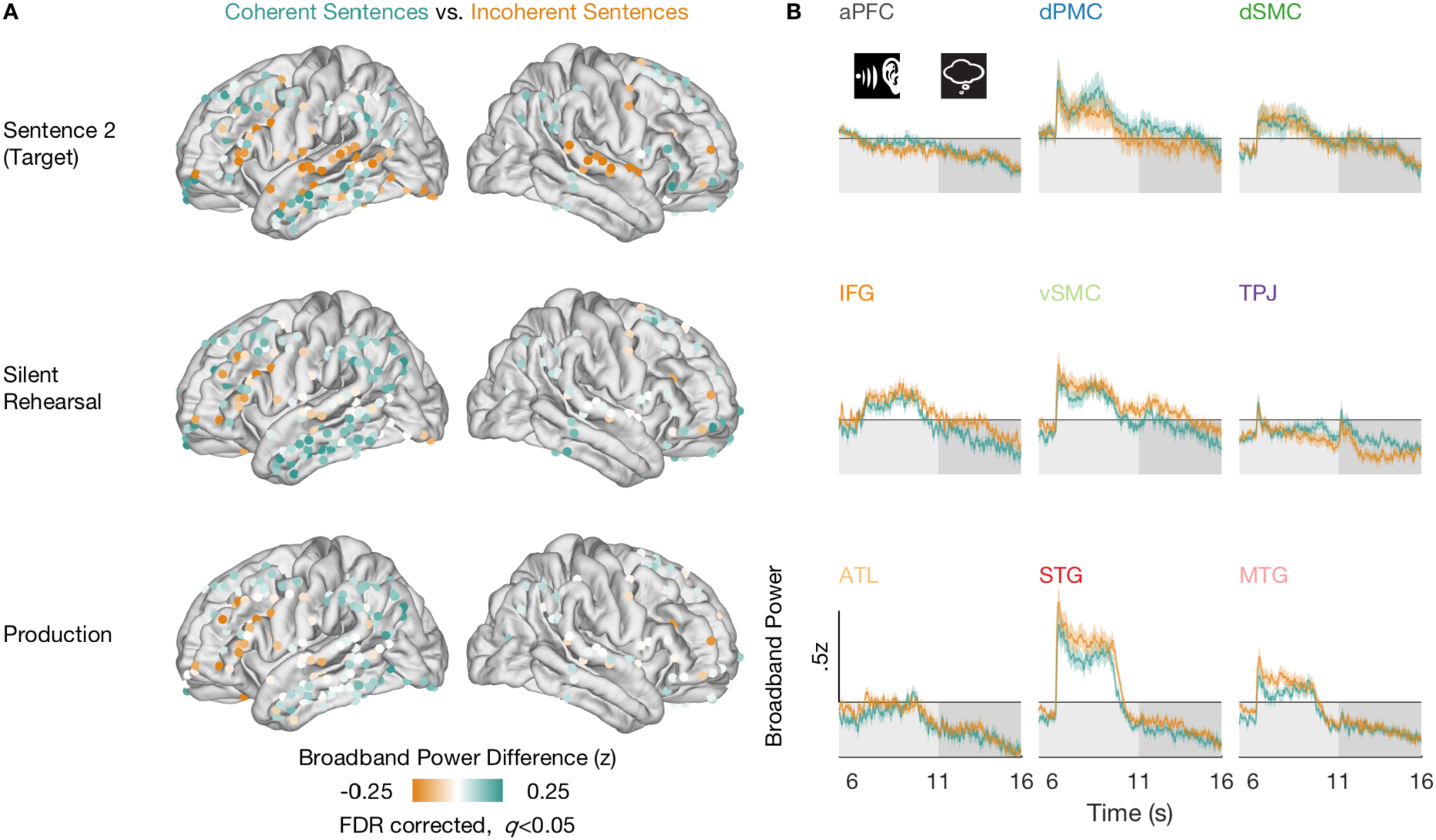
Perception and Silent Rehearsal of Coherent Sentences Produces Increased Broadband Power in Semantic Network. **A**, Spatial distribution of broadband power differences (coherent – incoherent) for coherent (teal) and incoherent sentences (orange) across task phases. Depicted are all sites showing a difference during either sentence 2, silent rehearsal or production. Only electrodes activated at the level *q* < 0.05 after FDR correction are shown. See also Figure S2. **B**, Average time courses for coherent and incoherent trials in the nine ROIs for the S2 and Rehearsal phases (indicated by icons and shading).

During perception, silent rehearsal and production of S2, coherent sentences were associated with greater broadband power in the middle and inferior temporal gyrus, temporoparietal cortex and in dorsal frontal areas (Figure 4A). However, overall activity in these areas was below baseline, suggesting that incoherent sentences were the cause of the reduction, while activity for coherent sentences was closer to baseline levels (Figure 4B). The pattern of greater broadband power for coherent sentences in posterior parietal, middle temporal and dorsal prefrontal cortex strongly resembles the “semantic network” identified by a meta-analysis of 120 fMRI studies (Binder et al., 2009).

During S2, perception of incoherent sentences elicited increased broadband power in left and right superior temporal gyri and left inferior frontal gyrus. The activation difference in the superior temporal gyrus disappeared during silent rehearsal and production but remained for the inferior frontal gyrus in those phases. Inspection of the time courses (Figure 4B) revealed that processing incoherent sentences resulted in stronger broadband power in all areas. This pattern of activation difference between coherent and incoherent sentences was stronger in the left hemisphere, though the overall pattern was similar in both hemispheres.

### Activation Time Courses are Sentence-Specific During Perception and Silent Rehearsal

Next, we assessed which of the widespread regions implicated in sentence perception and silent rehearsal were encoding sentence-specific information. A region might increase or decrease its activity due to general task demands (e.g., “listen”), without encoding information about the specific sentence that is being heard. Therefore, we measured whether activity time courses in each region were sentence-specific. Since we were interested in the nature of the representation during verbal short-term memory, we focused on sentence-specific representations during perception and silent rehearsal. To this end, we applied a technique for measuring region-level response reliability (Figure 5; see Methods, Reliability Analysis) to quantify sentence-specific responses in nine anatomical ROIs (Figure 2C). In each ROI, and separately for coherent and incoherent sentences, we subsampled stimuli to estimate a set of electrode weights (i.e., a spatial filter) that maximizes the reliability of responses across stimulus repeats (Figure 5A). Then, to obtain an unbiased estimate of how much sentence-specific information was contained in the time courses of each ROI, we measured the reliability of the weighted responses during the same task phase, but in different (out-of-sample) sentences and their repeats (Figure 5B).

**Figure 5.**
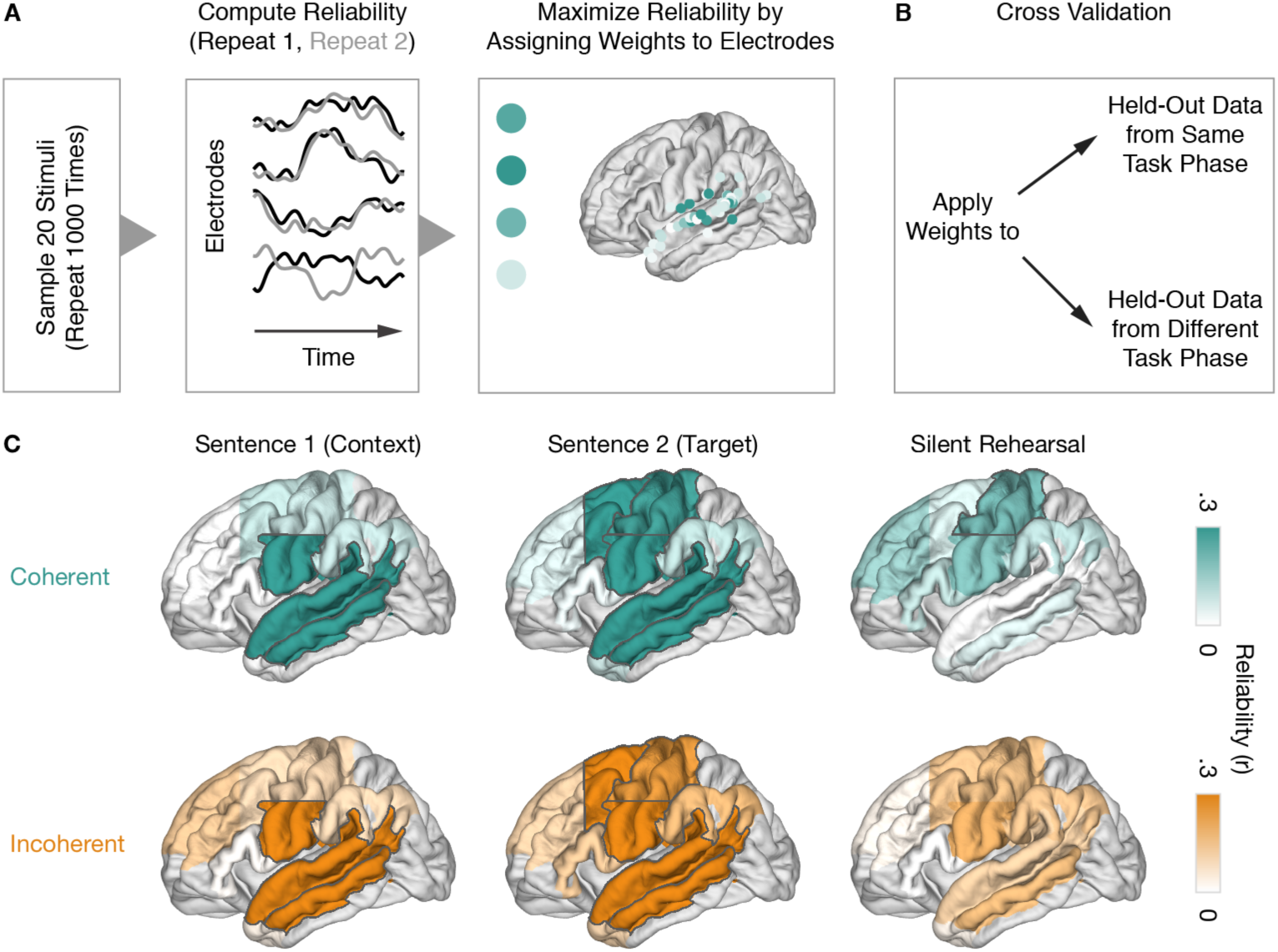
Sentence-Specific Activity in Sensory and Motor Cortices. Weighted reliability analysis workflow and stimulus-specific encoding during perception and silent rehearsal. **A**, Feature selection based on a subset of data. Reliability is maximized in a ROI by assigning different weights to electrodes. Data from eight participants who completed all trials went into this analysis. **B**, Cross validation using the weights from A on out-of-sample data from the same task phase (Figures 5, 6, S4A, S5, S6) or from a different task phase (Figures 6, 7, 8, S4B, S4C, S6). **C**, Sentence-specific encoding for coherent (teal, top) and incoherent sentences (orange, bottom) during perception (sentence 1, sentence 2) and silent rehearsal by ROI. The highlighted areas exhibit a significant difference from the null distribution (*p* < 0.05, two-sided, uncorrected). See also Figure S3 for electrode-level reliability. See also Figures S4A and S5A.

During perception, we observed that auditory, linguistic, and motor cortices of the left hemisphere exhibited the most reliable and sentence-specific responses to both coherent and incoherent sentences (Figures 5C, S5A; see Figure S3 for electrode-level reliability). During the auditory presentation of S1 (which did not have to be rehearsed), sentence-specific response time courses were observed in the left STG and MTG as well as in left vSMC. During the presentation of S2 (the target sentence for rehearsal), we additionally observed sentence-specific information in left dSMC and in left dPMC. These effects were observed for both coherent and incoherent sentences. In the right hemisphere (Figures S4A, S5B), sentence-specific responses were observed in the STG both during S1 and S2 and for coherent and incoherent sentences. Sentence-specific information was also present in the right vSMC, but this effect achieved statistical significance only for the coherent sentences.

During silent rehearsal, we observed reliable sentence-specific responses only in the left dSMC. This effect only achieved statistical significance for the coherent sentences but was similar for incoherent sentences. Looking more broadly across the surrounding areas, however, we did observe many weakly reliable responses across many temporal and peri-Rolandic regions during rehearsal (Figure S5). The within-phase reliability analysis could not determine whether these weakly reliable responses were truly sentence-specific, but we return to these areas in more powerful cross-phase analyses that have access to a larger sample of held-out sentences.

### Temporal Precision of Sentence-Specific Neural Activity

The previous analysis revealed sentence-specific neural activity that was most robust in temporal, sensorimotor and premotor cortices – but is this sentence-specific activity also specific to individual moments within the stimulus? To answer this question, we compared the sentence-specific activity across all pairs of individual time points during perception and during rehearsal. Using the weights optimized for each ROI in each fold of our cross-validation procedure, we computed the correlation of a specific moment’s activity level in the first presentation of a stimulus, comparing it against all possible timepoints in the second presentation of that stimulus.

The time-specific correlation analysis produces a two-dimensional time-time correlation matrix (left hemisphere, Figure 6; right hemisphere, Figure S6). The diagonal of the matrix reveals correlation at matching timepoints (time *t* in repeat 1 matched against the same point in repeat 2). If sentence-specific activity in a region varies from moment to moment, then the time-correlation values along the diagonal of the matrix will be higher than the off-diagonal values. Conversely, if the sentence-specific activity pattern evolves more slowly and is not specific to individual moments in the stimulus, then the temporal pattern will be more stationary, and non-matching timepoints will also be correlated across repeats; in this case, off-diagonal values will be as high as the on-diagonal.

**Figure 6.**
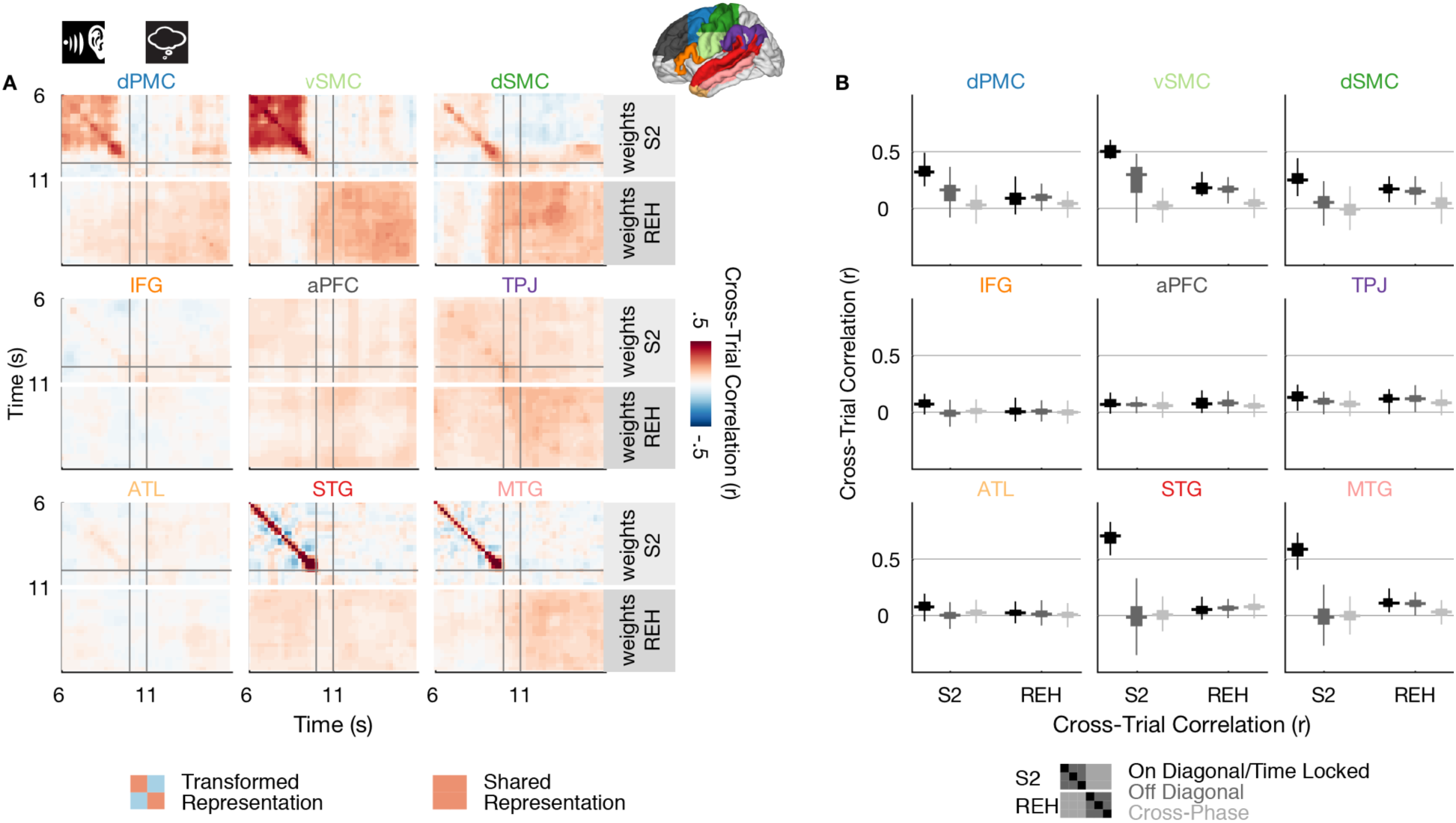
Temporal Specificity of Sentence-Locked Activity. See also Figure S6. **A**, Cross-time cross-trial correlation maps during perception and silent rehearsal. Within each ROI, a positive correlation at time (t1, t2) indicates that higher (spatially-weighted) activation values at time t1 on the first presentation of a sentence were associated with higher activation values at time t2 on the second presentation of the sentence. Thus, a positive correlation along the diagonal (t1=t2) indicates that responses are not only sentence-specific but also locked to individual timepoints within a stimulus. The correlations are computed either using the region-specific weights derived from the sentence 2 phase (S2, top part of each panel) or the weights derived from the rehearsal phase (REH, bottom part of each panel). Correlation values are averaged across coherent and incoherent sentences. The gray lines indicate the onsets of the silent gap following sentence 2 and the silent rehearsal cue, respectively. **B**, Statistical summary of the cross-time cross-trial correlation values shown in Panel A. The distribution of correlation values is shown for the on-diagonal component (t1=t2, black), the off-diagonal component (dark gray) and the cross-phase component (light gray) of the correlation matrix. Results are summarized separately for the sentence 2 and silent rehearsal phases and for each ROI. The fat horizontal line represents the median of the out-of-sample correlation distribution, while the box width represents its interquartile range. The vertical thin lines extend to the minimum/maximum of the out-of-sample correlation distribution, excluding outliers.

In the left hemisphere, we observed three sentence-specific response patterns, with varying degrees of temporal specificity (Figure 6). First, the STG and MTG exhibited sentence-locked activity only along the time-time diagonal and only during sentence perception. Thus, their responses are locked to specific moments within a sentence during perception, in line with their role in tracking low-level perceptual features (Cheung et al., 2016). A second pattern was observed in aPFC and TPJ: here the sentence-specific responses were temporally diffuse (not restricted to the time-time diagonal). Moreover, the reliable activity pattern was shared across perception and silent rehearsal; this pattern is suggestive of a slowly evolving representation that spans perception and rehearsal.

Finally, in vSMC, dSMC and dPMC, we observed a temporal encoding pattern that indicated a transformation between task phases. These sensorimotor circuits exhibited robust time-locked sentence-specific activity during sentence perception (i.e., elevated along the diagonal), combined with off-diagonal, slowly varying activity. During rehearsal, sentence-specific activity was more temporally diffuse. Notably, the sentence-specific activity in sensory and motor regions was distinct between perception and rehearsal phases: the electrode weights from perception showed little generalization to rehearsal (Figure 6B, cross-phase).

Among areas of the right hemisphere, only the STG exhibited sentence-locked activity along the diagonal during perception, suggesting a close tracking of auditory and phonological information as in the left hemisphere (Figure S6).

### Shared Sets of Electrodes in Prefrontal Cortex Sensitive to Contextual Information of the Past

The temporal specificity analyses (Figures 6, S6) suggest that, as we hear and then mentally rehearse sentences, some brain regions represent sentence-specific information in common sets of electrodes across the perceptual and rehearsal phases. To more quantitatively test this phenomenon, we applied our cross-validation procedure to the rehearsal phase using weights that were identified during perception of S2. Since the perceptual and rehearsal phases are completely separate data, this also enabled us to increase the power of our cross-validation procedure: we used the average weights estimated on 20 stimuli from the perceptual phase (Figures 5C, S4A) and measured the repeat correlations in the full set of 30 (instead of 10) held-out trials from the rehearsal phase. Using this approach, we identified sentence-specific activity with shared electrode weights in aPFC and TPJ. This cross-phase effect was observed separately for both coherent and incoherent sentences (Figure 7A). Thus, even though the sentence-specific activity in these high-level areas is weak and temporally diffuse (Figure 6), it is expressed in a common set of electrodes across perception of S2 and silent rehearsal.

**Figure 7.**
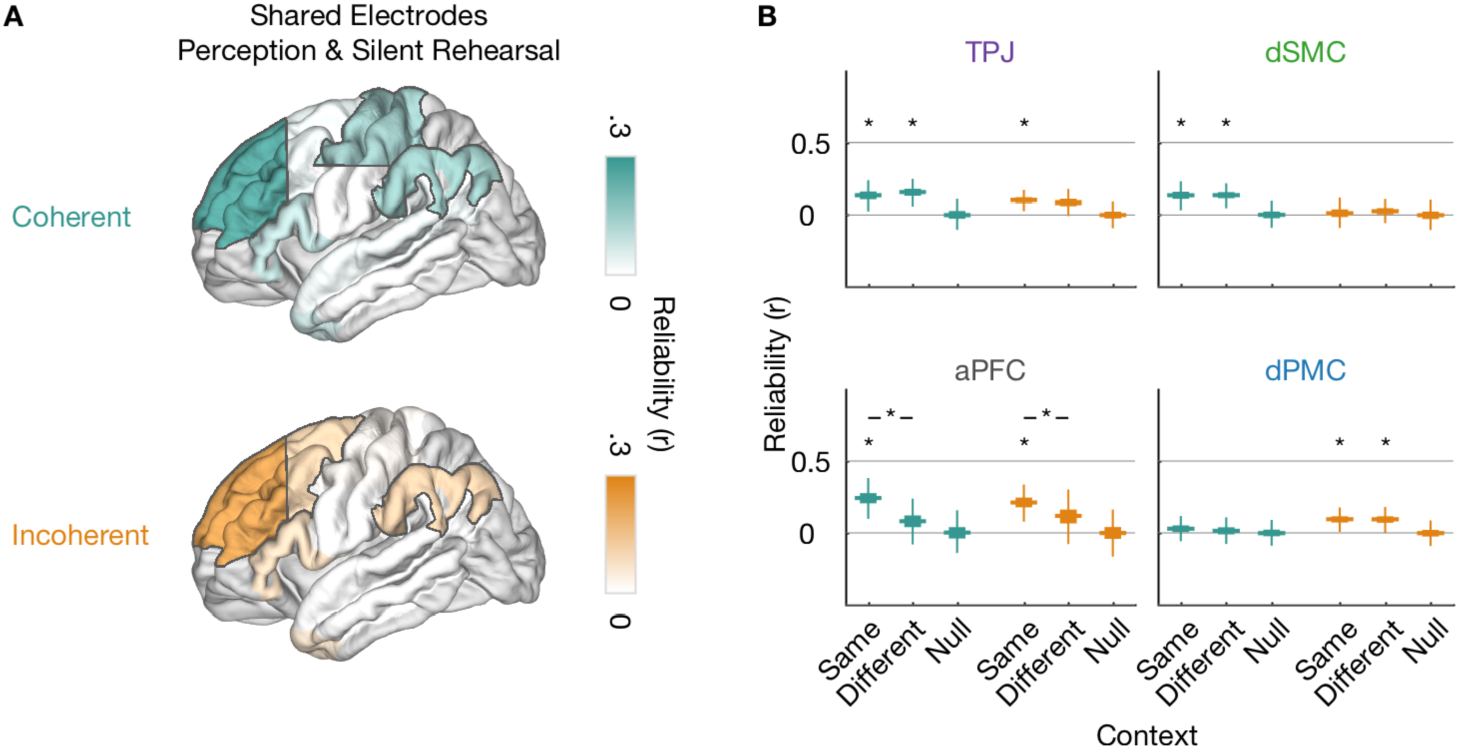
Shared Sets of Electrodes Encode Sentence-Specific Information. **A**, Reliability for shared electrodes in perception (sentence 2) and silent rehearsal that encode sentence-specific information for coherent (teal, top) and incoherent (orange, bottom) sentences. Weights from sentence 2 were applied to time courses from silent rehearsal. The highlighted areas exhibit a significant difference from the null distribution (*p* < 0.05, two-sided, uncorrected). See also Figure S4B. **B**, Reliability scores for the four ROIs sharing sets of electrodes between sentence 2 and silent rehearsal. Reliability is shown for the three cases when the second repeat of sentence 2 was preceded by the same sentence 1 context (Same) or the different sentence 1 context from the same stimulus group (Different). As a control, we also computed reliability for when both sentence 1 and sentence 2 were drawn from a nonmatching stimulus group (Null). Differences from the null distribution are indicated by single asterisks (*p* < 0.05 two-sided, uncorrected), differences between same vs. different contexts are indicated by lines (*p* < 0.05 one-sided, uncorrected).

We also observed significant sentence-specific information with shared electrode weights in dSMC (for coherent sentences only) and in dPMC (for incoherent sentences only). No sentence-specific activity with shared electrode weights was observed in the right hemisphere (Figure S4B).

What kind of information is encoded in the frontal and parietal electrodes that encode sentence-specific activity in both phases? Some models of verbal short-term memory have suggested that higher level sentence features (semantics and syntax) are used to “regenerate” information patterns at the time of sentence recall (Lombardi and Potter, 1992; Potter and Lombardi, 1990). Therefore, we tested whether the shared sentence-specific representations in each ROI were sensitive to the high-level semantics of the stimuli. To do so, we measured whether the sentence-specific activity in each region during silent rehearsal was the same across the two different semantic contexts that were generated by the non-rehearsed S1. For example, we tested whether the “spinning the wheel” sentence was represented the same way in the “ship captain” context and in the “game show” context” (see Figure 1B for both contexts). Thus, the phonological and syntactic features of the rehearsed sentence were identical, while its high-level semantics were altered by the context manipulation.

Only the aPFC was sensitive to the contextual information of S1 at the time of rehearsal (*p* < 0.05, one-sided, Figure 7B). In all other ROIs reliability was not detectibly affected by the contextual information of S1 (Figure 7B). This suggests that the most robustly encoded sentence-specific information during silent rehearsal (i.e., motor and premotor regions) were not representing high-level sentence context. Instead, the high-level information was present in the more temporally diffuse representations of the aPFC.

### Sentence-Specific Activity is Shared Between Perception and Rehearsal in Prefrontal and Temporoparietal Cortex

Given that common sets of electrodes encoded sentence-specific information across perception of S2 and silent rehearsal, we next assessed whether the sentence-specific neural activity patterns themselves were shared across perception and rehearsal. To this end, we directly compared the weighted and concatenated response time courses from the perception phase (S2) against the silent rehearsal phase. We employed the electrode weights from the S2 phase, and computed correlations against held-out data from a separate repeat: i.e., S2 phase of repeat 1 compared with silent rehearsal of repeat 2, and vice versa. We observed that sentence-specific time courses in aPFC, dPMC and TPJ of the left hemisphere were shared between perception and silent rehearsal. This effect was observed independently for coherent and incoherent sentences (Figure 8). For coherent sentences, sentence-specific time courses were also shared across task phases in the vSMC. In the right hemisphere, sentence-specific time courses were shared only in the MTG and only for incoherent sentences (Figure S4C).

**Figure 8.**
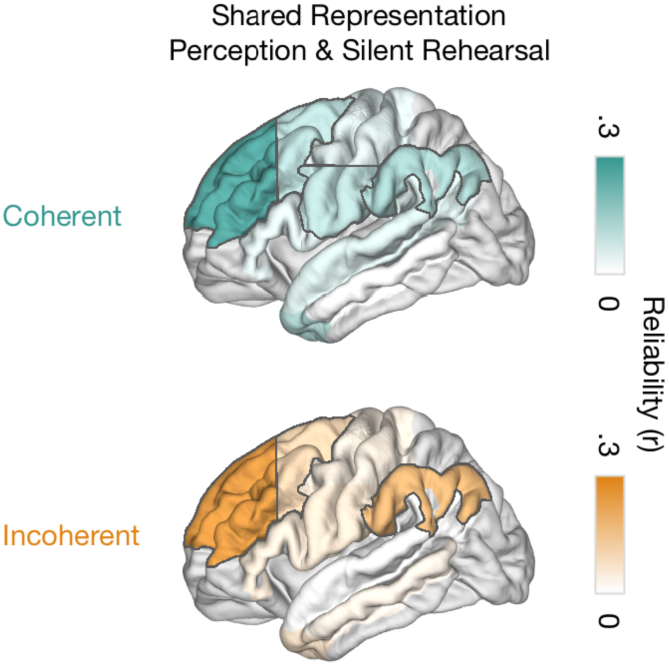
Shared Sentence-Specific Time Courses Between Perception and Silent Rehearsal. Sentence-specific reliability measured by directly correlating the response during perception (sentence 2) and during silent rehearsal. Cross-phase correlations for coherent (top) and incoherent (bottom) sentences are shown in the left hemisphere. Weights for this cross-phase analysis were defined as those that maximized the within-phase reliability in sentence 2. The highlighted areas exhibit a significant difference from the null distribution (*p* < 0.05, two-sided, uncorrected). See also Figure S4C.

## Discussion

We recorded intracranial neural activity across widespread language-related networks as people heard and then mentally rehearsed spoken sentences. In each brain region, we tested whether silent rehearsal of sentences involved reactivation of sentence-specific representations established during perception, or transformation to a distinct sentence-specific representation. We found evidence for both processes: transformation was most apparent in SMC and PMC, while aPFC and TPJ maintained a more static representation that was shared across phases. In the aPFC and TPJ, where representations were shared across perception and rehearsal, neural representation were also more sensitive to changes in the syntactic structure and contextual semantics of the sentences.

The data implicate sensorimotor cortex (both vSMC and dSMC) and PMC as critical bridges between the perception and rehearsal of spoken sentences. These regions were the only ones for which neuronal activity, as indexed by broadband power, was consistently increased during both silent rehearsal and perception (Figure 3A). Moreover, broadband power in motor and premotor areas during rehearsal predicted the accuracy of sentence memory (Figure 3B). Finally, vSMC, dSMC and PMC exhibited stimulus-specific encoding across both perceptual and silent rehearsal phases (Figure 5) and this stimulus-specific encoding during perception was temporally precise (Figure 6). These findings are consistent with prior ECoG reports of short-latency encoding of auditory speech in SMC and PMC (Cheung et al., 2016; Edwards et al., 2009; Glanz et al., 2018), and they extend the prior work by tracking the representations of complex speech in motor areas during the transition from perception to rehearsal.

Sentence-specific activity in vSMC, dSMC and PMC was transformed between the perception and silent rehearsal of the same sentence. A moment-by-moment comparison of the sentence-specific activity (Figure 6) revealed a transition between distinct representations in perception and silent rehearsal. Sentence-specific activity patterns defined during perception in the vSMC, dSMC and PMC did not extend beyond the end of the perception of S2; at the same time, a distinct rehearsal-specific activity pattern became reliable, starting from the end of the perception phase and continuing into rehearsal. Thus, the present data connect SMC and PMC to a transformation process that supports short-term memory for natural spoken language.

These findings are consistent with a sensorimotor transformation model that has been proposed for working memory (Cogan et al., 2014, 2017); they are not consistent with models that posit a one-to-one mirroring of activity during perception and production (D’Ausilio et al., 2009; see also Liberman et al., 1967).

Short-term memory for phonological information is thought to be supported by an auditory-motor interface (Buchsbaum and D’Esposito, 2008; Hickok and Poeppel, 2007; Jacquemot and Scott, 2006; Rauschecker, 2011). The present data indicate that the SMC and PMC could possess both the auditory and motor representations necessary for such an interface. The vSMC activity in speech perception likely reflects more low-level and obligatory audio-motor representations (Cheung et al., 2016): this area exhibited reliable sentence-specific activity even for S1 (which did not have to be rehearsed), and its reliability increased only marginally for S2 (which had to be rehearsed). By contrast, the dSMC and PMC were more sensitive to the task relevance of the speech input: these areas exhibited greater sentence-specific reliability for S2 (which had to be rehearsed), relative to S1 (Figures 5C, S5). We tentatively propose that the vSMC representations directly track auditory and motor representations, while the time-locked dSMC and PMC responses are purposed not only toward control of articulatory sequences, but also toward expectation of sensory sequences (e.g., Schubotz and von Cramon, 2003). A short-term memory trace of speech in sensorimotor cortices could also explain why these areas would be engaged for discriminating speech stimuli in noise (Du et al., 2014).

In contrast with the sensorimotor circuits discussed above, the prefrontal and temporoparietal cortex exhibited a temporally diffuse sentence-specific activity pattern, which was shared across the perception and silent rehearsal of the same sentence. Although some of the sentence-specific activity in motor sites was also shared across phases (Figures 7,8), these shared effects were inconsistent across conditions and made up only a small proportion of the reliable signal in motor circuits (Figure 6B). By contrast, the cross-phase correlation in aPFC and in the TPJ was as large as the within-phase correlations (Figure 6B, within-phase and cross-phase). At the level of aggregate activation, the prefrontal and temporoparietal circuits exhibited increased broadband power responses for coherent sentences (Figure 4), consistent with their role in a “semantic network” (Binder et al., 2009).

Prefrontal and temporoparietal cortex have long temporal receptive windows, exhibiting slower population dynamics than sensory cortices (Honey et al., 2012; Murray et al., 2014) and responding to new input in a manner that depends on prior context (Hasson et al., 2015; Lerner et al., 2011). Consistent with this prediction, the temporal activation pattern in the left aPFC was sensitive to prior contextual information (changes in the high-level situational meaning of S2 based on the context provided by S1). The stable sentence-specific signals in higher order areas may provide a persistent “scaffold” which supports “regeneration” of detailed surface features of the sentence (e.g., phonemes and prosody) from more abstract properties (e.g., syntax and semantics) (Lombardi and Potter, 1992; Potter and Lombardi, 1990; see also Savill et al., 2017). More generally, the slowly changing prefrontal representations we observed in high level cortical areas are consistent with a distributed, drifting cortical “context” that recursively interacts with episodic retrieval processes (Polyn et al., 2009). We note, however, that the temporal context effects we observed were small relative to the those in prior studies using naturalistic stimuli (e.g., Lerner et al., 2011); rich and extended narrative stimuli likely generate a more powerful contextual modulation than the single preceding sentence (S1) that we used in the present design.

Sentence-specific information changed most rapidly in the posterior areas of frontal cortex (i.e. in motor cortex) and changed more slowly toward anterior regions (i.e. premotor and prefrontal cortex, Figure 6). The distinct timescales of these frontal areas parallel the recent observation of distinct working memory functions for populations of neurons with distinct dynamical timescales (Cavanagh et al., 2018; Wasmuht et al., 2018). Prefrontal neurons with intrinsically short timescales, as measured by spontaneous firing dynamics, responded more rapidly during the perceptual phase of a task, whereas those with intrinsically longer timescales encoded information in a more stable way towards a delay period (Wasmuht et al., 2018). In our task, faster dynamics seem to be more important in sensorimotor transformation (from perception to subvocal rehearsal), whereas slower dynamics were associated with areas whose function may be more semantically sensitive. In future work, we will directly characterize the relative timing across these different stages of processing, in order to determine changes in the direction of information flow across perception and rehearsal.

Unexpectedly, coherent sentences and incoherent sentences were represented in a stimulus-specific manner within essentially the same sets of brain regions, albeit with different levels of mean broadband power activation. This suggests that even the incoherent sentences possessed sufficient high-level structure (e.g., lexical semantics and aspects of syntax), similar to that used to represent the coherent sentences. This minor effect of sentence incoherence is consistent with the general observation that semantic implausibility has less of an effect on sentence processing than strong syntactic violations (Polišenská et al., 2015).

During silent rehearsal, we expected to observe, but did not observe, sentence-specific activity in the ATL (Patterson et al., 2007), the posterior STG and planum temporale (Buchsbaum and D’Esposito, 2008), and the IFG (Hickok, 2012). There is evidence of marginally reliable responses during the perception phase in both the ATL and in the IFG in the time-resolved analysis (Figure 6B). In addition, one electrode in the posterior temporal cortex was jointly active in both perception and rehearsal, but we did not find stimulus-specific decoding in the superior temporal cortex as a whole during silent rehearsal. It is possible that stimulus-specific encoding could be identified in these areas given a larger pool of patients, denser electrode coverage, or depth electrode coverage targeting sulci (especially the Sylvian fissure). Although we had broad coverage, we cannot rule out the possibility of false negatives; instead, our data speaks most clearly for the reliable and stimulus-specific responses across the lateral cerebral cortex, spanning perception and rehearsal.

Overall, our data suggest that sensorimotor and premotor cortices can support an audio-motor interface proposed by leading models of verbal short-term memory. In the SMC and PMC we observed extensive joint activation across perception and rehearsal, and rehearsal activity which predicted the accuracy of later sentence production. In parallel with this possible audio-motor interface, more abstract sentence features were maintained in prefrontal and temporoparietal circuits. To better understand how high-level semantic features constrain and facilitate our inner speech, future work should examine the interactions between sensorimotor circuits and the frontal and temporoparietal cortex as we silently rehearse sequences of words.

## Acknowledgments

We thank the patients for participating in this study and the staff of the epilepsy monitoring at Toronto Western Hospital. We thank Victoria Barkley for assisting with data collection and David Groppe for support with the iElvis toolbox. We are grateful for funding support from the DFG (MU 3875/2-1 to K.M.), the National Institute of Mental Health (R01 MH111439-01, subaward C.J.H), the Sloan Foundation (Research Fellowship to C.J.H.), the National Science Foundation (NSF DMS-1811315 to K.M.T.), and the Toronto General and Toronto Western Hospital Foundation.

## Supplement

**Figure S1.**
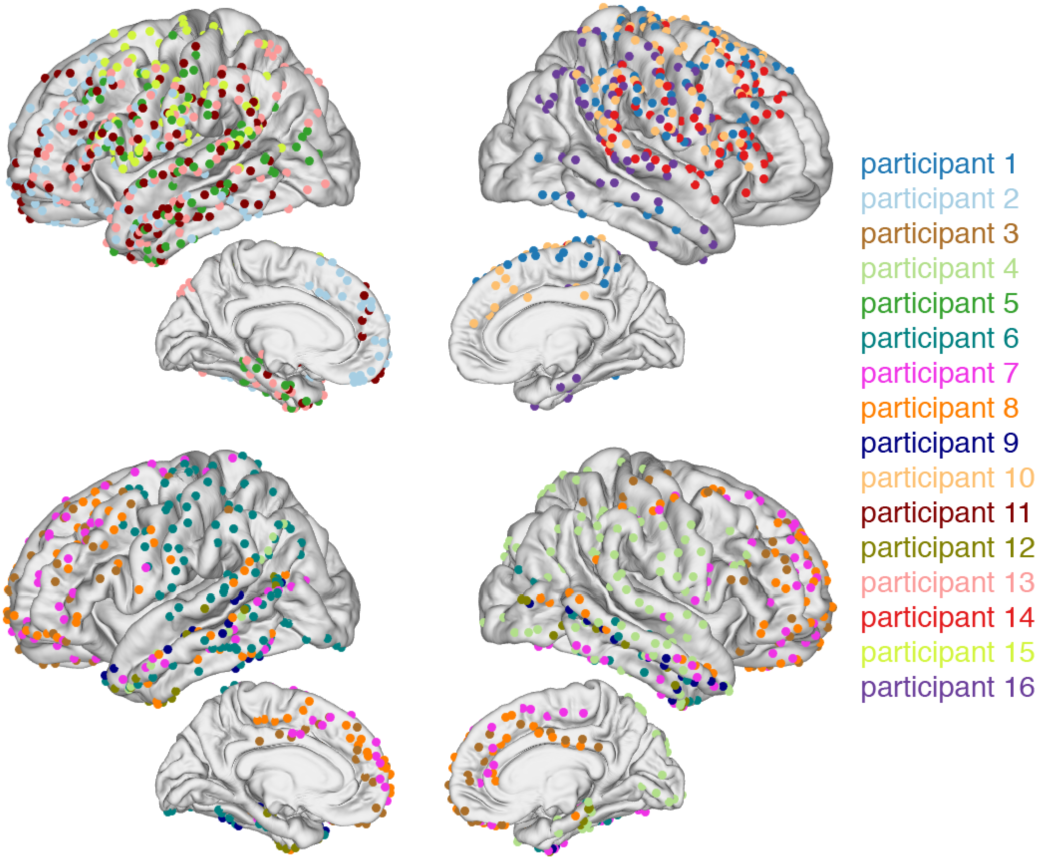
Electrode Placement for Individual Participants Coded by Color. Related to Figure 1D.

**Figure S2.**
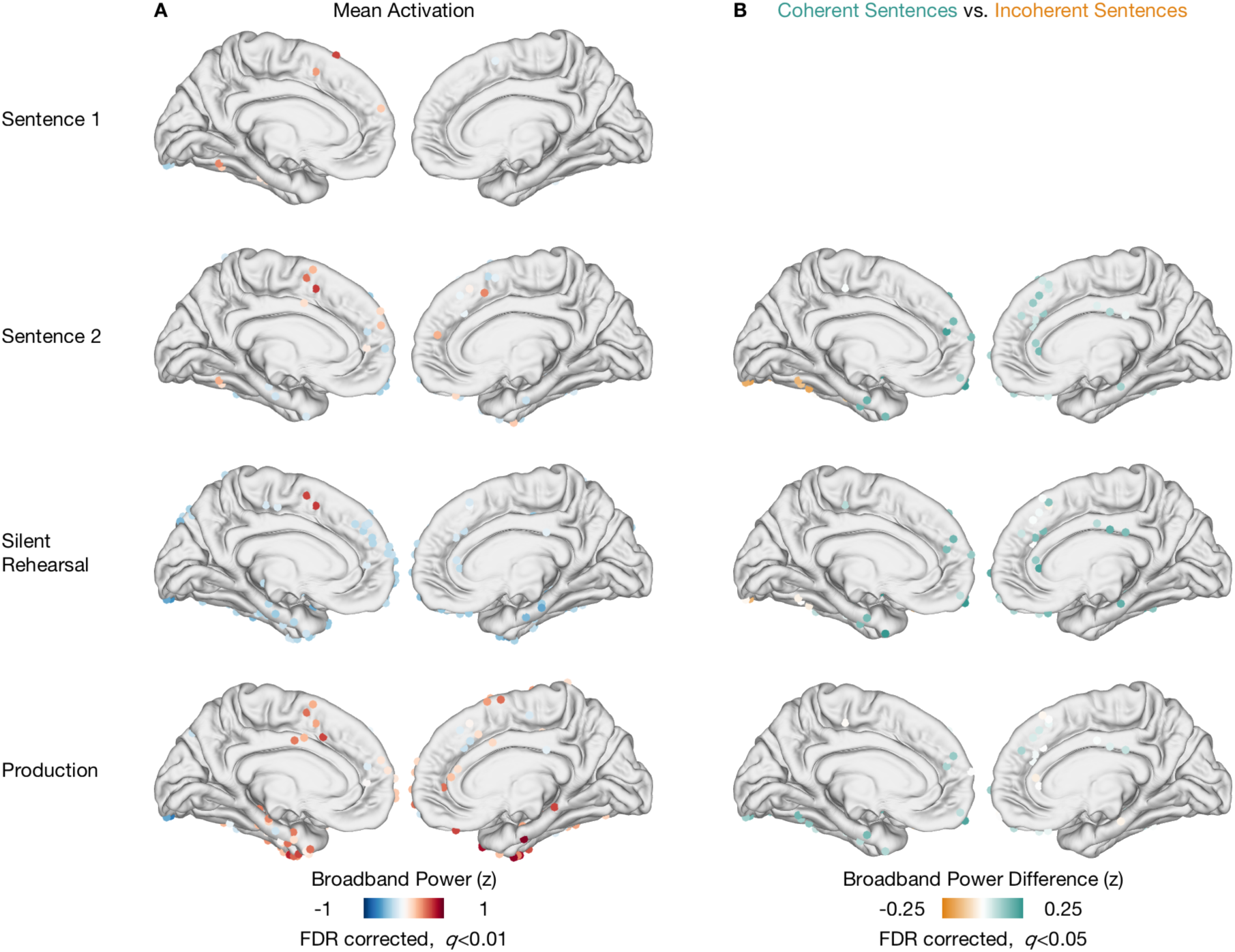
Spatial Distributions of Broadband Power Responses in Medial Cortex. Related to Figures 2B and 4A. **A**, Spatial distribution of broadband power responses aggregated across participants and conditions for each task phase in the left and right hemispheres. Only electrodes activated at the level q < 0.01 after FDR correction are shown. **B**, Spatial distribution of broadband power differences between coherent and incoherent sentences across task phases. Depicted are all sites showing a difference during either sentence 2, silent rehearsal or production. Only electrodes activated at the level q < 0.05 after FDR correction are shown.

**Figure S3.**
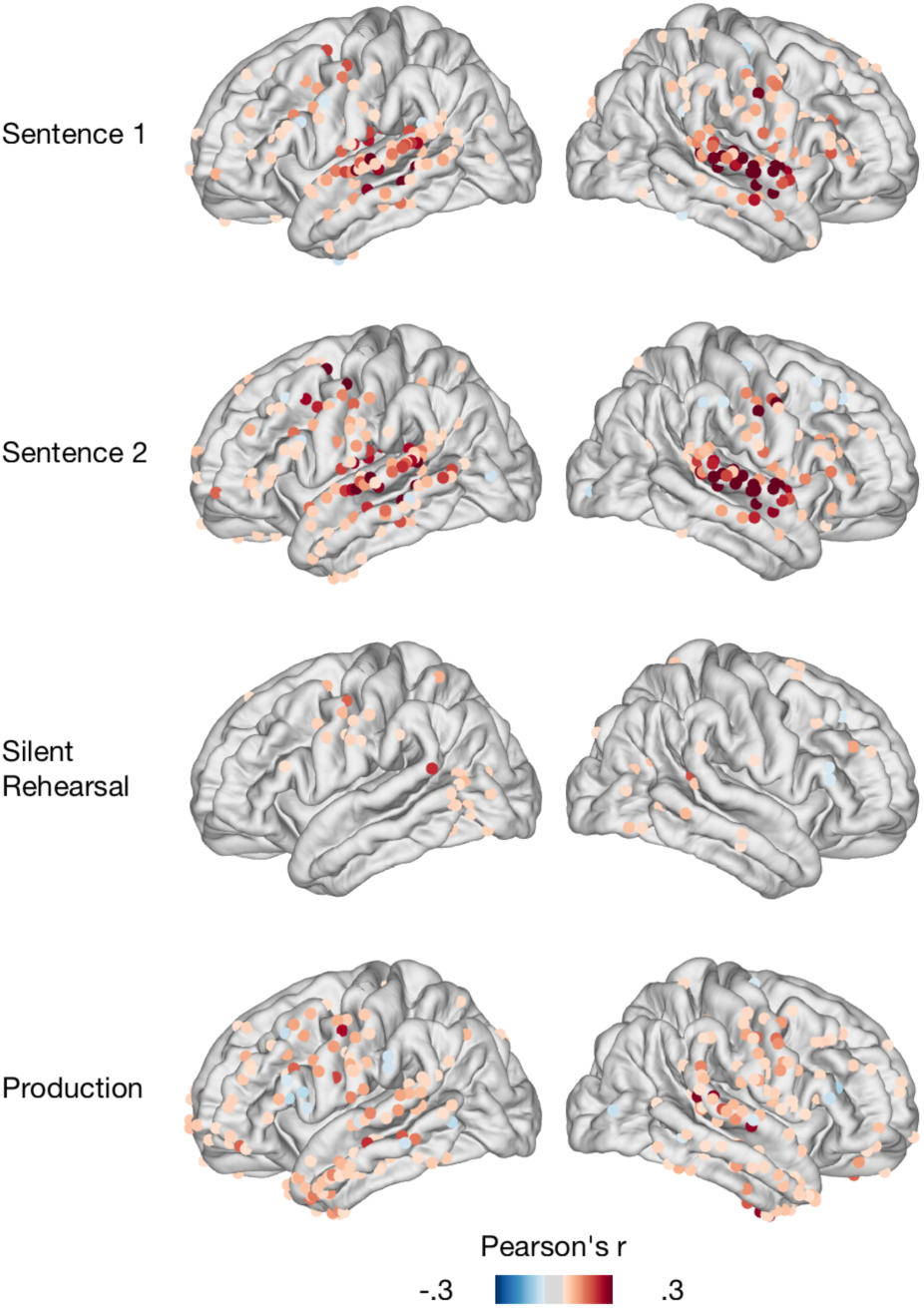
Spatial Distributions of Average Within-Trial Repeat Reliability in All Task Phases. Related to Figures 5C and S4A. Pearson’s *r* correlation values were computed separately for each trial and its repeat and then averaged. For perception (sentence 1, sentence 2), the first and last 500 ms were discarded to avoid on- and offset effects on reliability. For production, the first 1000 ms was discarded to avoid onset effects. Only *r* values smaller than −0.05 and exceeding 0.05 are shown.

**Figure S4.**
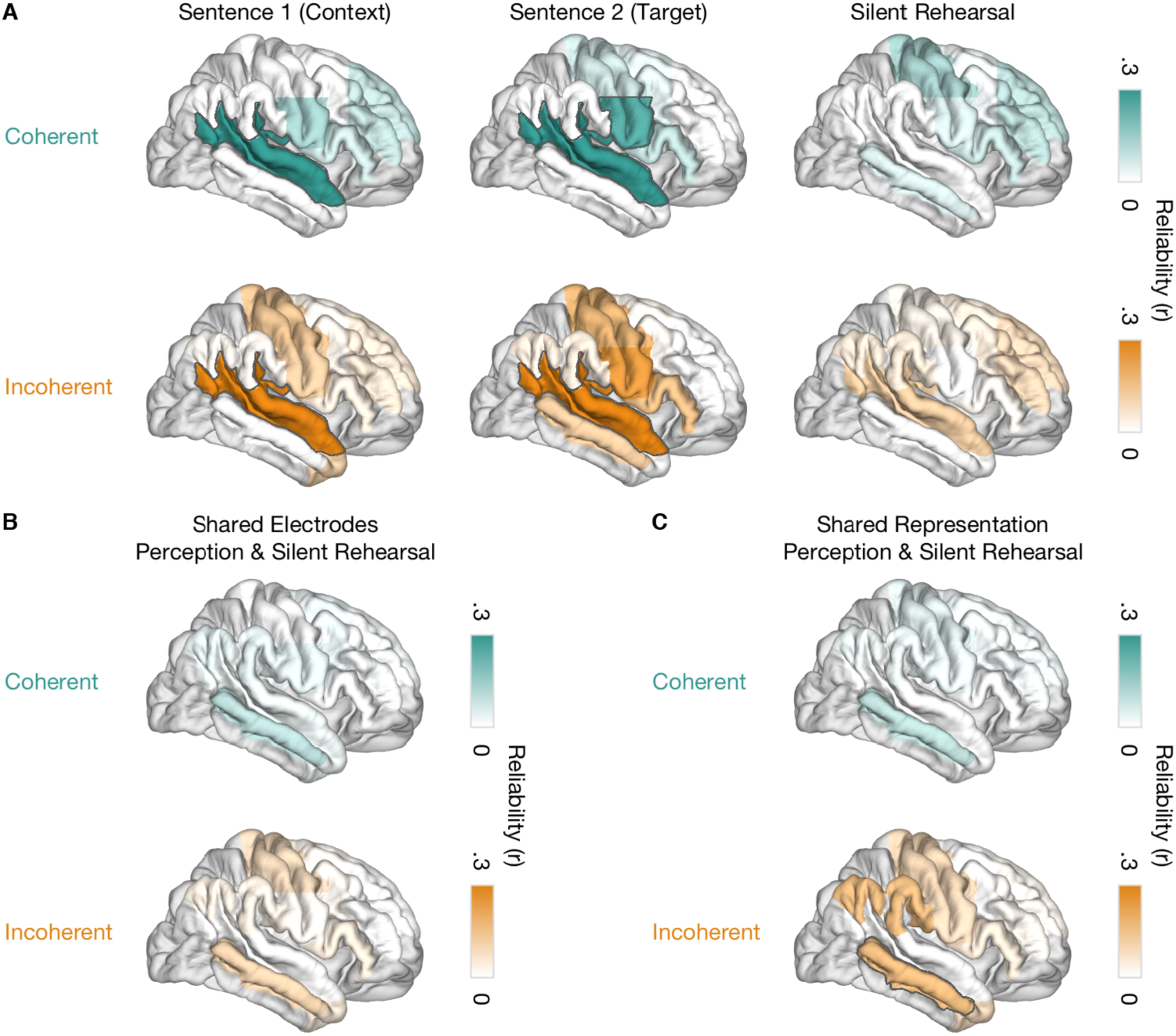
Results of the Correlated Component Analysis in the Right Hemisphere. Related to Figures 5C, 7A and 8. **A**, Sentence-specific encoding in the right hemisphere during perception (sentence 1, sentence 2) and silent rehearsal. Reliability is shown for each ROI for coherent (top) and incoherent sentences (bottom). The highlighted areas exhibit a significant difference from the null distribution (*p* < 0.05, two-sided, uncorrected). See also Figures S3 and S5B. **B**, Reliability for shared electrodes in the right hemisphere. No ROI shares electrodes encoding sentence-specific information for coherent (top) and incoherent (bottom) sentences during perception (sentence 2) and silent rehearsal. Weights from sentence 2 were applied to time courses from silent rehearsal. **C**, Sentence-specific reliability directly correlating perception (sentence 2) and silent rehearsal for coherent (top) and incoherent (bottom) sentences. Weights from sentence 2 were used for feature selection. The highlighted area exhibits a significant difference from the null distribution (*p* < 0.05, two-sided, uncorrected).

**Figure S5.**
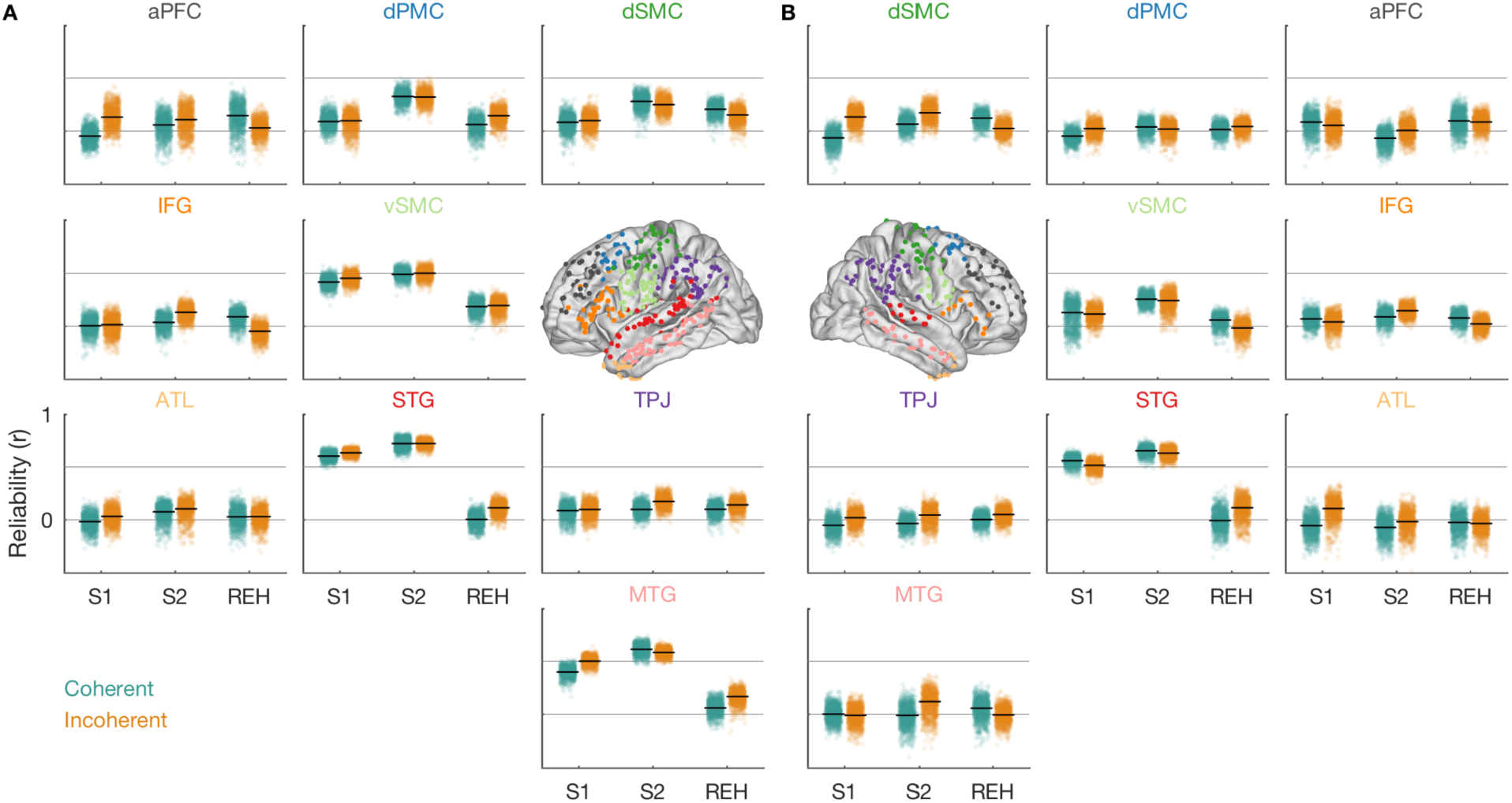
Stimulus-Specific Encoding During Sentence 1 (S1), Sentence 2 (S2) and Silent Rehearsal (REH). Related to Figures 5C and S4A. Shown is the reliability for each of the 1,000 permutations in the nine ROIs in the **A**, left and **B**, right hemispheres. The horizontal black line indicates the mean across all permutations, which is identical to the reliability shown in Figures 5 and S4A.

**Figure S6.**
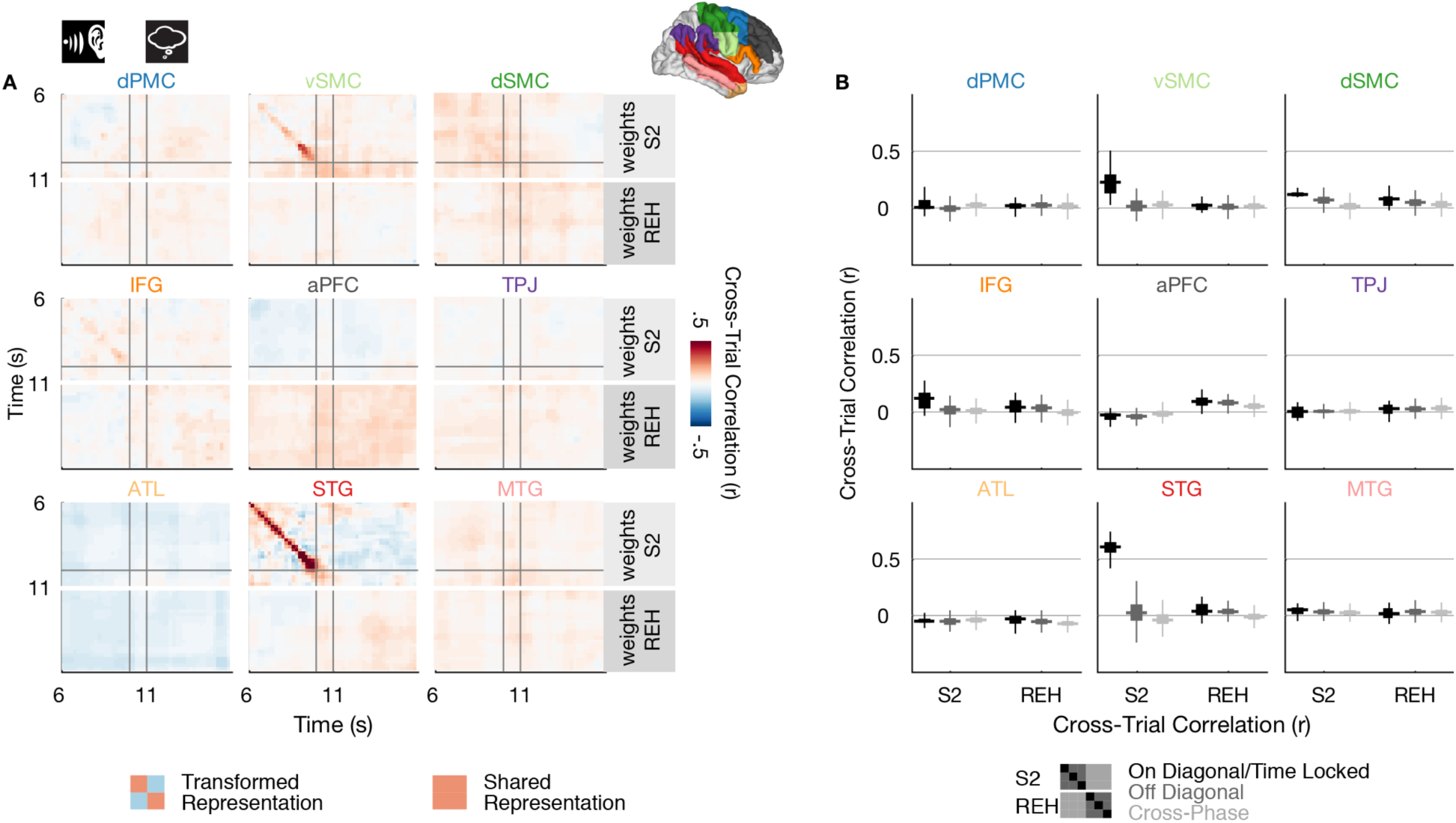
Temporal Specificity of Sentence-Locked Activity in the Right Hemisphere. Related to Figure 6. **A**, Cross-time cross-trial correlation maps during perception and silent rehearsal. Within each ROI, a positive correlation at time (t1, t2) indicates that higher (spatially-weighted) activation values at time t1 on the first presentation of a sentence were associated with higher activation values at time t2 on the second presentation of the sentence. Thus, a positive correlation along the diagonal (t1=t2) indicates that responses are not only sentence-specific but also locked to individual timepoints within a stimulus. The correlations are computed either using the region-specific weights derived from the sentence 2 phase (S2, top part of each panel) or the weights derived from the rehearsal phase (REH, bottom part of each panel). Correlation values are averaged across coherent and incoherent sentences. The gray lines indicate the onsets of the silent gap following sentence 2 and the silent rehearsal cue, respectively. **B**, Statistical summary of the cross-time cross-trial correlation values shown in Panel A. The distribution of correlation values is shown for the on-diagonal component (t1=t2, black), the off-diagonal component (dark gray) and the cross-phase component (light gray) of the correlation matrix. Results are summarized separately for the sentence 2 and silent rehearsal phases and for each ROI. The fat horizontal line represents the median of the out-of-sample correlation distribution, while the box width represents its interquartile range. The vertical thin lines extend to the minimum/maximum of the out-of-sample correlation distribution, excluding outliers.

**Table S1.** **Clinical and Demographic Information for the Participating Patients.** Related to Methods, Participants.

| Participant | Gender | Age at Surgery/<br>Seizure Onset | Language<br>Laterality | Language (Age of<br>Onset) | Seizure<br>Type | Seizure Focus |
| --- | --- | --- | --- | --- | --- | --- |
| P1 | F | 50.7/8 | L | English | CPS | L Post Mesial |
| P2 | M | 28.4/16 | L | English | CPS | Multifocal |
| P3 | F | 24.8/12 | R | English | CPS | L Fronto-Temporal |
| P4 | F | 28.7/17 | L | English | CPS | Bilateral<br>Occipital |
| P5 | F | 26.2/19 | NA | English | CPS | L Inferior<br>Temporal |
| P6 | F | 32.4/18 | Bilateral (R>L) | English (0), Greek<br>(0) | CPS | Unclear |
| P7 | M | 28.0/1.5 | NA | English | CPS | Unclear |
| P8 | M | 36.9/14 | L | English | Deja-vu,<br>GCS | L (poorly<br>localized) |
| P9 | M | 35.2/4 | Unclear | English | CPS | Bilateral<br>Temporal |
| P10 | F | 25.1/5 | L | Portuguese (0),<br>English (3) | CPS | Diffuse |
| P11 | F | 19.7/<1 | Bilateral (L>R) | Rumanian (0),<br>English (<13) | CPS | L Temporal |
| P12 | F | 45.4/25 | L | English | CPS | R Anterior<br>Temporal |
| P13 | F | 39.4/19 | Bilateral (R>L) | English | CPS, GTS | Unclear |
| P14 | F | 18.8/12 | NA | English | CPS | R Frontal |
| P15 | M | 28.1/7 | L | English | NA | Multifocal |
| P16 | F | 28.0/21 | L | English | CPS,<br>GTCS | R Temporal |

*Note:* CPS, complex partial seizure; F, female; GCS, generalized clonic seizure; GTCS, generalized tonic clonic seizure; GTS, generalized tonic seizure; L, left; M, male; NA, not available; R, right.

**Table S2.**
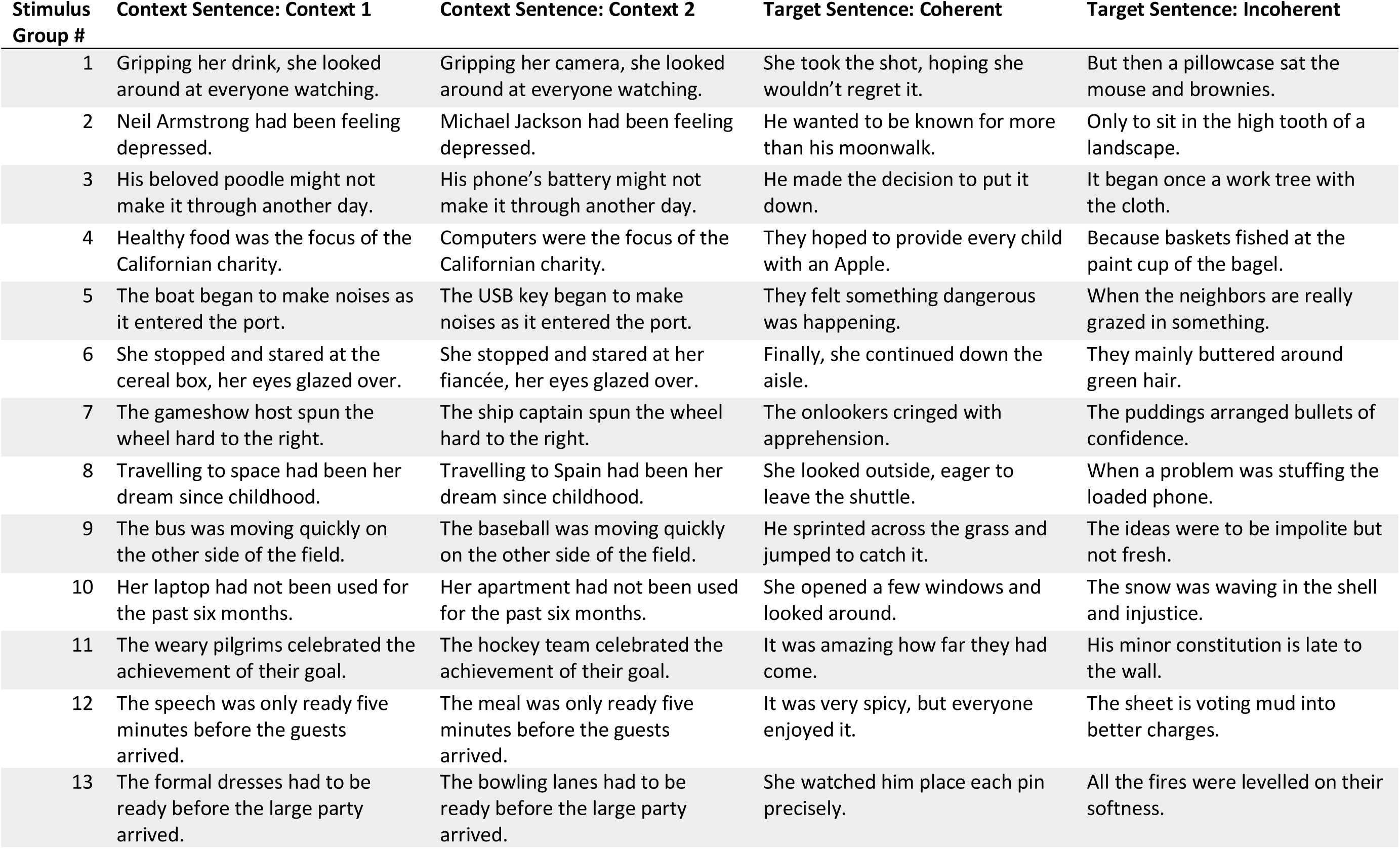
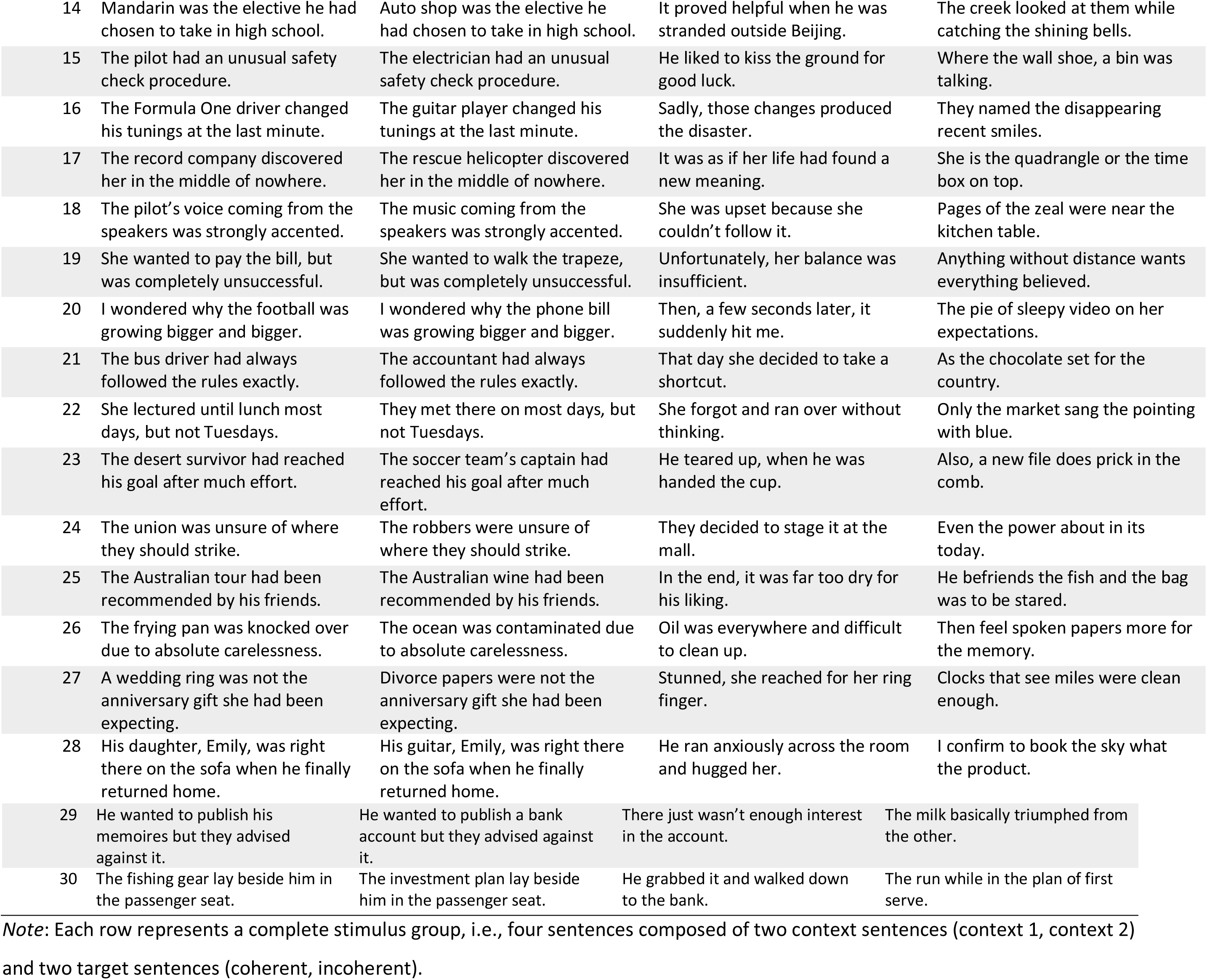
**Full Sentence List.** Related to Figure 1B.

| Stimulus Group # | Context Sentence: Context 1 | Context Sentence: Context 2 | Target Sentence: Coherent | Target Sentence: Incoherent |
| --- | --- | --- | --- | --- |
| 1 | Gripping her drink, she looked around at everyone watching. | Gripping her camera, she looked around at everyone watching. | She took the shot, hoping she wouldn't regret it. | But then a pillowcase sat the mouse and brownies. |
| 2 | Neil Armstrong had been feeling depressed. | Michael Jackson had been feeling depressed. | He wanted to be known for more than his moonwalk. | Only to sit in the high tooth of a landscape. |
| 3 | His beloved poodle might not make it through another day. | His phone's battery might not make it through another day. | He made the decision to put it down. | It began once a work tree with the cloth. |
| 4 | Healthy food was the focus of the Californian charity. | Computers were the focus of the Californian charity. | They hoped to provide every child with an Apple. | Because baskets fished at the paint cup of the bagel. |
| 5 | The boat began to make noises as it entered the port. | The USB key began to make noises as it entered the port. | They felt something dangerous was happening. | When the neighbors are really grazed in something. |
| 6 | She stopped and stared at the cereal box, her eyes glazed over. | She stopped and stared at her fiancée, her eyes glazed over. | Finally, she continued down the aisle. | They mainly buttered around green hair. |
| 7 | The gameshow host spun the wheel hard to the right. | The ship captain spun the wheel hard to the right. | The onlookers cringed with apprehension. | The puddings arranged bullets of confidence. |
| 8 | Travelling to space had been her dream since childhood. | Travelling to Spain had been her dream since childhood. | She looked outside, eager to leave the shuttle. | When a problem was stuffing the loaded phone. |
| 9 | The bus was moving quickly on the other side of the field. | The baseball was moving quickly on the other side of the field. | He sprinted across the grass and jumped to catch it. | The ideas were to be impolite but not fresh. |
| 10 | Her laptop had not been used for the past six months. | Her apartment had not been used for the past six months. | She opened a few windows and looked around. | The snow was waving in the shell and injustice. |
| 11 | The weary pilgrims celebrated the achievement of their goal. | The hockey team celebrated the achievement of their goal. | It was amazing how far they had come. | His minor constitution is late to the wall. |
| 12 | The speech was only ready five minutes before the guests arrived. | The meal was only ready five minutes before the guests arrived. | It was very spicy, but everyone enjoyed it. | The sheet is voting mud into better charges. |
| 13 | The formal dresses had to be ready before the large party arrived. | The bowling lanes had to be ready before the large party arrived. | She watched him place each pin precisely. | All the fires were levelled on their softness. |
| 14 | Mandarin was the elective he had chosen to take in high school. | Auto shop was the elective he had chosen to take in high school. | It proved helpful when he was stranded outside Beijing. | The creek looked at them while catching the shining bells. |
| 15 | The pilot had an unusual safety check procedure. | The electrician had an unusual safety check procedure. | He liked to kiss the ground for good luck. | Where the wall shoe, a bin was talking. |
| 16 | The Formula One driver changed his tunings at the last minute. | The guitar player changed his tunings at the last minute. | Sadly, those changes produced the disaster. | They named the disappearing recent smiles. |
| 17 | The record company discovered her in the middle of nowhere. | The rescue helicopter discovered her in the middle of nowhere. | It was as if her life had found a new meaning. | She is the quadrangle or the time box on top. |
| 18 | The pilot's voice coming from the speakers was strongly accented. | The music coming from the speakers was strongly accented. | She was upset because she couldn't follow it. | Pages of the zeal were near the kitchen table. |
| 19 | She wanted to pay the bill, but was completely unsuccessful. | She wanted to walk the trapeze, but was completely unsuccessful. | Unfortunately, her balance was insufficient. | Anything without distance wants everything believed. |
| 20 | I wondered why the football was growing bigger and bigger. | I wondered why the phone bill was growing bigger and bigger. | Then, a few seconds later, it suddenly hit me. | The pie of sleepy video on her expectations. |
| 21 | The bus driver had always followed the rules exactly. | The accountant had always followed the rules exactly. | That day she decided to take a shortcut. | As the chocolate set for the country. |
| 22 | She lectured until lunch most days, but not Tuesdays. | They met there on most days, but not Tuesdays. | She forgot and ran over without thinking. | Only the market sang the pointing with blue. |
| 23 | The desert survivor had reached his goal after much effort. | The soccer team's captain had reached his goal after much effort. | He teared up, when he was handed the cup. | Also, a new file does prick in the comb. |
| 24 | The union was unsure of where they should strike. | The robbers were unsure of where they should strike. | They decided to stage it at the mall. | Even the power about in its today. |
| 25 | The Australian tour had been recommended by his friends. | The Australian wine had been recommended by his friends. | In the end, it was far too dry for his liking. | He befriends the fish and the bag was to be stared. |
| 26 | The frying pan was knocked over due to absolute carelessness. | The ocean was contaminated due to absolute carelessness. | Oil was everywhere and difficult to clean up. | Then feel spoken papers more for the memory. |
| 27 | A wedding ring was not the anniversary gift she had been expecting. | Divorce papers were not the anniversary gift she had been expecting. | Stunned, she reached for her ring finger. | Clocks that see miles were clean enough. |
| 28 | His daughter, Emily, was right there on the sofa when he finally returned home. | His guitar, Emily, was right there on the sofa when he finally returned home. | He ran anxiously across the room and hugged her. | I confirm to book the sky what the product. |
| 29 | He wanted to publish his memoirs but they advised against it. | He wanted to publish a bank account but they advised against it. | There just wasn't enough interest in the account. | The milk basically triumphed from the other. |
| 30 | The fishing gear lay beside him in the passenger seat. | The investment plan lay beside him in the passenger seat. | He grabbed it and walked down to the bank. | The run while in the plan of first serve. |

*Note:* Each row represents a complete stimulus group, i.e., four sentences composed of two context sentences (context 1, context 2) and two target sentences (coherent, incoherent).

**Table S3.**
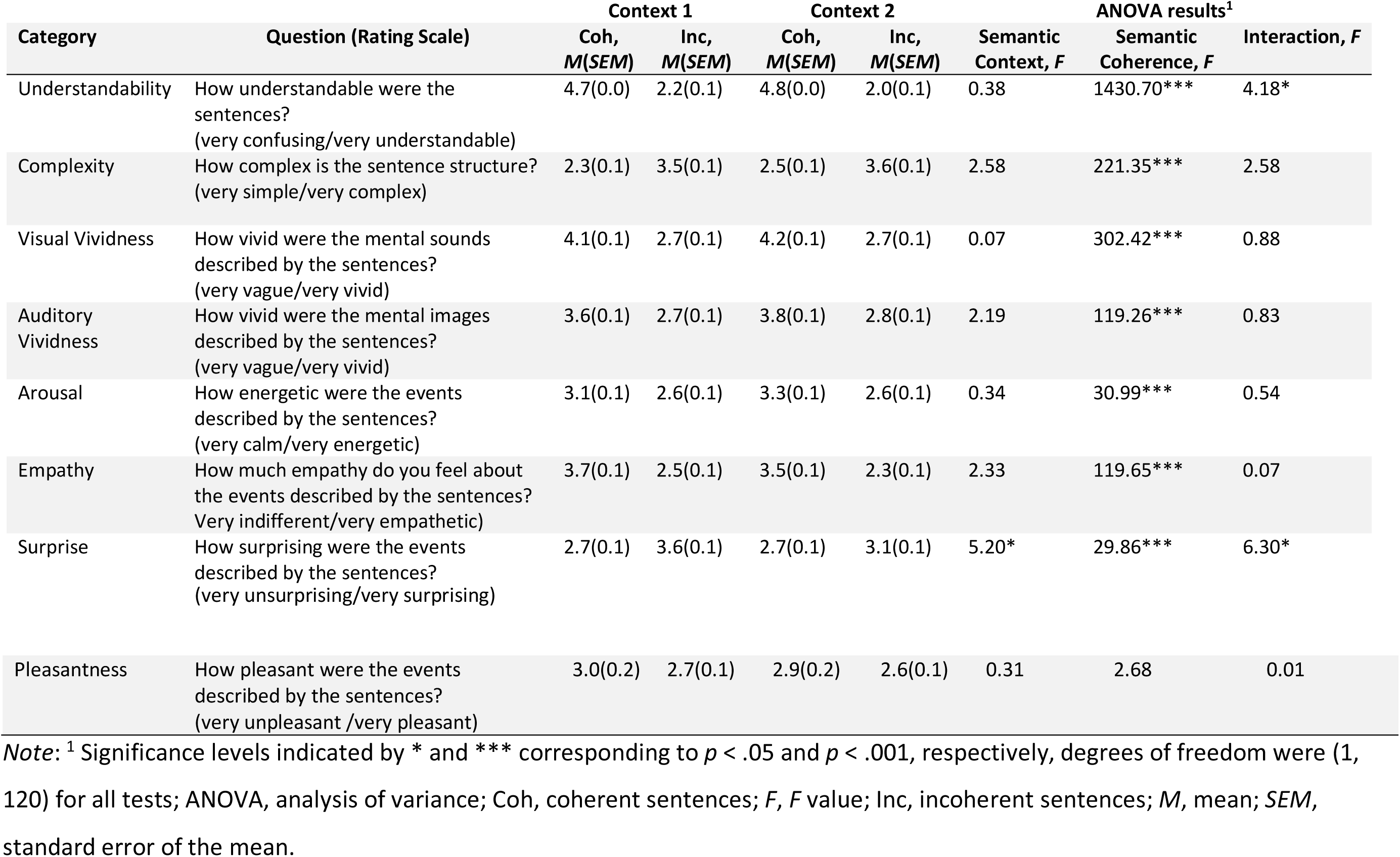
**Questions Used in and Results of Norming Task.** Related to Figure 1C.

**Table S4.** **Included Areas in Destrieux Atlas per Region of Interest (ROI).** Related to Figure 2C.

| ROI | List of Included Areas in Destrieux Atlas |
| --- | --- |
| Superior Temporal Gyrus (STG) | G_temp_sup-Lateral, G_temp_sup-Plan_polar, G_temp_sup-Plan_tempo, Lat_Fis-post, S_temporal_sup |
| Middle Temporal Gyrus (MTG) | G_temporal_middle,S_temporal_inf |
| Anterior Temporal Lobe (ATL) | Pole_temporal |
| Temporoparietal Junction (TPJ) | G_pariet_inf-Angular, G_pariet_inf-Supramar |
| Inferior Frontal Gyrus | G_front_inf-Opercular, G_front_inf-Orbital, G_front_inf-Triangul |
| Ventral Sensory Motor Cortex (vSMC) | G_and_S_paracentral, G_and_S_subcentral, G_postcentral, G_precentral, S_central, S_postcentral, S_precentral-inf-part, S_precentral-sup-part |
| Dorsal Sensory Motor Cortex (dSMC) | G_and_S_paracentral, G_and_S_subcentral, G_postcentral, G_precentral, S_central, S_postcentral, S_precentral-inf-part, S_precentral-sup-part |
| Dorsal Premotor Cortex (dPMC) | G_front_middle, G_front_sup, S_front_inf, S_front_middle, S_front_sup |
| Anterior Prefrontal Cortex (aPFC) | G_front_middle, G_front_sup, S_front_inf, S_front_middle, S_front_sup |

**Author contributions** (CRediT taxonomy)
Conceptualization, K.M.and C.J.H.; Methodology, K.M. and C.J.H.; Formal Analysis, K.M., K.M.T., and C.J.H.; Investigation, K.M. and K.H.; Resources, K.M., K.M.T., T.A.V., and C.J.H.; Writing – Original Draft, K.M., K.H., K.M.T., T.A.V., and C.J.H.; Visualization, K.M., K.M.T., and C.J.H.; Supervision, T.A.V. and C.J.H.; Funding Acquisition, K.M., K.M.T., T.A.V. and C.J.H.

